# Dynamics of the cell fate specifications during female gametophyte development in *Arabidopsis*

**DOI:** 10.1101/2020.04.07.023028

**Authors:** Daichi Susaki, Takamasa Suzuki, Daisuke Maruyama, Minako Ueda, Tetsuya Higashiyama, Daisuke Kurihara

**Affiliations:** Kihara Institute for Biological Research, Yokohama City University, Yokohama, Japan; Department of Biological Chemistry, College of Bioscience and Biotechnology, Chubu University, Kasugai, Japan; Institute of Transformative Bio-Molecules (ITbM), Nagoya University, Nagoya, Japan; Division of Biological Science, Graduate School of Science, Nagoya University, Nagoya, Japan; Department of Biological Sciences, Graduate School of Science, University of Tokyo, Tokyo, Japan; JST, PRESTO, Nagoya, Japan

**Keywords:** Arabidopsis, cell fate specification, female gametophyte, live-cell imaging, transcriptome

## Abstract

The female gametophytes of angiosperms contain cells with distinct functions, such as those that enable reproduction via pollen tube attraction and fertilization. Although the female gametophyte undergoes unique developmental processes, such as several rounds of nuclear division without cell plate formation, and the final cellularization, it remains unknown when and how the cell fate is determined during their development. Here, we visualized the living dynamics of female gametophyte development and performed transcriptome analysis of its individual cell types, to assess the cell fate specifications in *Arabidopsis thaliana*. We recorded time lapses of the nuclear dynamics and cell plate formation from the one-nucleate stage to the seven-cell stage after cellularization, using the *in vitro* ovule culture system. The movies showed that the nuclear division occurred along the micropylar–chalazal axis. During cellularization, the polar nuclei migrated while associating with forming edge of the cell plate. Then, each polar nucleus migrated to fuse linearly towards each other. We also tracked the gene expression dynamics and identified that the expression of the *MYB98pro∷GFP*, a synergid-specific marker, was initiated before cellularization, and then restricted to the synergid cells after cellularization. This indicated that cell fates are determined immediately after cellularization. Transcriptome analysis of the female gametophyte cells of the wild type and *myb98* mutant, revealed that the *myb98* synergid cells had the egg cell-like gene expression profile. Although in the *myb98*, the egg cell-specific gene expressions were properly initiated only in the egg cells after cellularization, but subsequently expressed ectopically in one of the two synergid cells. These results, together with the various initiation timings of the egg cell-specific genes suggest the complex regulation of the individual gametophyte cells, such as cellularization-triggered fate initiation, MYB98-dependent fate maintenance, cell morphogenesis, and organelle positioning. Our system of live-cell imaging and cell-type-specific gene expression analysis provides insights into the dynamics and mechanisms of cell fate specifications in the development of female gametophytes in plants.

## 1 Introduction

In multicellular organisms, each differentiated cell creates complex structures to perform its specified functions. As cells differentiate according to their cell fate, it is important for the cell fate to be determined at the appropriate time and position. However, the molecular mechanisms that determine how cells recognize positional information and their cell fate in plants are not well understood. The development of the female gametophyte in angiosperms is of interest when studying cell fate specifications, as they are essential for cell differentiation in plants.

The female gametophytes in angiosperms contain highly differentiated cells with distinct functions, those for such as pollen tube attraction and fertilization, which enable plant reproduction. In *Arabidopsis thaliana*, one megaspore undergoes three rounds of mitosis without cytokinesis as a coenocyte. Cellularization occurs almost simultaneously around each nucleus, producing the *Polygonum*-type female gametophyte with eight nuclei and seven cells: one egg cell, one central cell, two synergid cells, and three antipodal cells. It is important for the sexual reproduction of angiosperms that each cell of the female gametophyte develops by acquiring its appropriate cell fate. Although it remains unknown when and how the cell fate is determined during female gametophyte development, two mechanisms are thought to play important roles: cell polarity along the micropyle–chalazal axis in the female gametophyte and cell–cell communications after cellularization. The female gametophytes of angiosperms develops with distinct polarity. In many plant species, the egg cell and the synergid cells form at the micropylar end of the ovule, and the antipodal cells form at the opposite side of the chalazal end (Maheshwari, 1950; Yadegari and Drews, 2004).

The egg cell is the intrinsic female gamete that forms the embryo in the seeds by fertilization with the sperm cell carried by the pollen tube. The central cell is the largest in the female gametophyte, and often contains multiple nuclei during cellularization. In the case of *Polygonum* type female gametophytes, the central cell contains two nuclei (Schmid et al., 2015). The central cell is regarded as one of the gametes because it fertilizes the sperm cell, but the fertilized central cell forms the embryo-nursing tissue endosperm in the seed, and it is not inherited in the next generation. The synergid cells have a morphology thought to be specialized for secretion. The synergid cell has finger-like plasma membrane invaginations with thickened cell walls termed “filiform apparatus” in the micropylar end. This structure increases the surface area of the synergid cells with a higher rate of exocytosis for secretion. When the pollen tube arrives at the synergid cells, the synergid cells stop elongation of the pollen tube and cause the release of sperm cells by rupturing its tip (Higashiyama, 2002).

These three cell types are highly common among angiosperm species and are essential for sexual reproduction. Except for the antipodal cells, the set of the egg cell, the central cell, and the synergid cells have been designated as “egg apparatus” that are essential for sexual reproduction (Huang and Russell, 1992). In contrast, the function of the antipodal cells is poorly understood and varies widely among plant species (Diboll and Larson, 1966; Maeda and Miyake, 1997; An and You, 2004; Holloway and Friedman, 2008; Heydlauff and Groß-Hardt, 2014). For example, it has been found that the antipodal cells degenerated by programmed cell death as the female gametophytes mature in *Arabidopsis* and *Torenia fournieri* (Yadegari and Drews, 2004; Mól, 1986). However, other report has suggested that antipodal cells did not degenerate but persisted beyond fertilization in *Arabidopsis* (Song et al., 2014).

Mutant screening and gene expression analysis are two major approaches to explore the factors responsible for their cell fates. Large-scale mutant screenings have been carried out with mutagenesis by T-DNA insertions or transposons, to identify the genes required for female gametophyte development (reviewed in Brukhin et al. 2005). For instance, *lachesis* and *eostre* were reported as the mutants whose synergid cell fates changed to egg cell-like (Groß-Hardt et al., 2007; Pagnussat et al., 2007). On the other hand, several reverse genetic investigations, based on gene expression analysis, reported the identification of important genes which had cell-type specific functions. First, the gene expression comparisons between the ovules, with or without the female gametophyte, identified *MYB98*, a synergid specific transcription factor in *Arabidopsis* (Kasahara et al., 2005). Further analysis, including *myb98* ovules compared to controls clarified the putative female gametophyte-specific gene cluster controlled by MYB98 (Yu et al., 2005; Jones-Rhoades et al., 2007; Steffen et al., 2007). The pollen tube attractants, ZmEA1 and TfLUREs, were identified in gene expression analysis of maize egg cells and the *Torenia* synergid cells (Márton et al., 2005; Okuda et al., 2009; Márton et al., 2012).

The technology and techniques for the gene expression analyses of the female gametophytes have advanced over time. Initially, RT-PCR-based screenings were performed with the ovules of the wild type or the female gametophyte mutants (Kasahara et al., 2005). Then, microarray analyses were developed for use with the ovules of a wild-type and the mutants without the female gametophyte or *myb98* in *Arabidopsis* (Yu et al., 2005; Jones-Rhoades et al., 2007; Steffen et al., 2007). Further detailed gene expression analyses in each type of cell have been reported, such as the expressed sequence tag analysis for the protoplasts of the maize and wheat egg cells and the *Torenia* synergid cells (Sprunck et al., 2005; Márton et al., 2005; Okuda et al., 2009). The protoplasts of rice egg and synergid cells and the *Arabidopsis* egg cells, synergid cells, and central cells, which were collected by laser-assisted microdissection (LAM), were analyzed with a microarray (Ohnishi et al., 2011; Wuest et al., 2010). These studies showed the genome-wide gene expression profiles of each cell-type in the rice and mature *Arabidopsis* ovules. In recent years, RNA-sequencing (RNA-seq) has become a major technology in transcriptomics. In plant reproduction research, the protoplasts of rice egg cells, sperm cells, and pollen vegetative cells and the protoplasts of *Arabidopsis* egg cells, zygotes in their early stages, embryos, and the central cells collected by LAM, have been investigated by RNA-seq (Anderson et al., 2013; Zhao et al., 2019; Schmid et al., 2012). These reports have demonstrated that RNA-seq could detect greater levels of gene expression than microarrays and the genome-wide gene expression profiles at higher resolutions. From these studies, the characteristics of each female gametophyte cells have been identified, and the genes responsible for each cells function have gradually been elucidated.

As described above, the two major methods of sample collection to analyze the female gametic transcriptome were LAM or manual isolation of the protoplasts. Protoplasts collection was technically challenging and had lower costs (Wuest et al., 2013) but was expected to extract more RNA as it used living cells. The protoplast isolation of female gametophyte cells was previously reported in many species (Theunis et al. 1991; *Torenia fournieri*, Mól 1986; *Plumbago zeylanica*, Huang and Russell 1989; *Zea maize*, Kranz et al. 1991; *Oryza sativa* Uchiumi et al. 2006). In most studies, female gametophyte cells were isolated with the enzyme solution containing cellulase. The optimized conditions of the enzyme solution were different for different plant species (Kawano et al., 2011). The cell-type specific RNA-seq, including the mutants defective in female gametophyte cell function, must be powerful tools to reveal the precise gene expression changes associated with each cell functions or specifications. Convenient and simple methods for cell isolation enabled these analyses.

Genes expressed specifically in each female gametophyte cell and used as markers of the cell fate have been identified in several plants, particularly *Arabidopsis* (Tekleyohans et al., 2017). However, it is not clear when and how these cells specify their cell fates and exhibit specific gene expressions. The mutant analysis shows a strict correlation between nuclear position and cell fate (Kong et al., 2015; Groß-Hardt et al., 2007; Pagnussat et al., 2007; Moll et al., 2008; Kirioukhova et al., 2011). It is still unknown whether the nuclear positions determine the cell fates or not, due to little spatio-temporal information on the detailed nuclear dynamics and cell fate specifications. As the female gametophyte development occurs deep within the female pistil, it has been challenging to observe directly in the living state. Therefore, the intracellular behavior of the female gametophyte development has been analyzed by fixing the ovules and observing the sections. It is crucial to capture the living dynamics in the female gametophyte development to reveal the dynamics of cell fate specification.

Here, we performed live-cell imaging of the female gametophytes development in *Arabidopsis* using the *in vitro* ovule culture system, which enabled us to observe the nuclear dynamics, division, cellularization, and cell fate specifications in real-time, by using specific fluorescent marker lines. Subsequently, we established a method for the isolation of each female gametophyte cells with high efficiency, without contaminating the other cells in *Arabidopsis*. We then built a technology platform for transcriptome analysis using a next-generation sequencer for a small number of isolated female gametophyte cells. Furthermore, we analyzed the contributions of the cell-cell communications in changing the gene expressions, by analyzing the expression profiles of the synergid cells of the *myb98* mutant, a transcription factor that is thought to contribute to the determination of the synergid cell fate.

## 2 MATERIALS AND METHODS

For all experiments, the *Arabidopsis thaliana* accession Columbia (Col-0) was used as the wild type. The following transgenic lines were previously described: *RPS5Apro∷H2B–tdTomato* (Adachi et al., 2011), *RPS5Apro∷tdTomato–LTI6b* (Mizuta et al., 2015), *RPS5Apro∷H2B–sGFP* (Maruyama et al., 2015), *FGR8.0* (Völz et al., 2013), *MYB98pro∷GFP* (Kasahara et al., 2005), *EC1.2pro∷mtKaede* (Hamamura et al., 2011), *FWApro∷FWA–GFP* (Kinoshita et al., 2004), and *ABI4pro∷H2B–tdTomato* (Kimata et al., 2016).

*Arabidopsis* seeds were sown on plates containing half-strength Murashige and Skoog salts (Duchefa Biochemie, Haarlem, The Netherlands), 0.05% MES-KOH (pH 5.8), 1× Gamborg’s vitamin solution (Sigma, St Louis, MO, USA), and 1% agar. The plates were incubated in a growth chamber at 22°C under continuous lighting after cold treatments at 4°C for 2—3 days in the dark. Two-week-old seedlings were transferred to soil and grown at 21 to 25°C under long-day conditions (16-h light/8-h dark).

### 2.1 Plasmid Construction

The *GPR1pro∷H2B–mNeonGreen* (coded as DKv1200), was constructed with the 2,568 bp upstream regions of *GPR1* (At3g23860) and the full-length coding region of *H2B* (HTB1: At1g07790), fused to the *mNeonGreen* (Allele Biotechnology, San Diego, CA) with the (SGGGG)_2_ linker, and the 1,959 bp downstream regions were cloned into the binary vector pPZP211 (Hajdukiewicz et al., 1994). The *CDR1–LIKE2pro∷CDR1–LIKE2–mClover* (coded as DKv1023) was constructed using the 1,398 bp upstream regions and the full-length coding region of *CDR1–LIKE2* (At1g31450), fused to the *mClover* with the (SGGGG)_2_ linker, and the *NOS* terminator, and cloned into the binary vector pPZP211. The *CDR1–LIKE1pro∷CDR1–LIKE1–mClover* (coded as DKv1024) was constructed using the 2,000 bp upstream regions and the full-length coding region of *CDR1–LIKE1* (At2g35615) fused to the *mClover* with the (SGGGG)_2_ linker and the *NOS* terminator, and then cloned into the binary vector pPZP211. Finally, the *CDR1pro∷CDR1–mClover* (coded as DKv1025) was constructed with the 1,577 bp upstream regions and the full-length coding region of *CDR1* (At5g33340) fused to the *mClover* with the (SGGGG)_2_ linker, and the *NOS* terminator, cloned into the binary vector pPZP211.

To construct the multiple cell-type-specific marker line with the nuclei marker (coded as DKv1110), the following sequences were cloned into the binary vector pPZP211 and the *NPTII* replaced with *mCherry* under the control of the *At2S3* promoter from a pAlligator-derived binary vector (Kawashima et al., 2014): *EC1.1pro∷SP–mTurquoise2–CTPP* (Kimata et al., 2019) (the 463 bp *EC1.1* promoter was fused to *mTurquoise2* that fused to the signal peptide (SP) sequence of *EXGT–A1* (At2g06850) at the N-terminus and to a vacuolar sorting signal COOH-terminal propeptide (CTPP), and the *HSP* terminator); *DD1pro∷ermTFP1* (the 1,262 bp *DD1* promoter (At1g36340) was fused to mTFP1 that was fused to the SP sequence of *EXGT–A1* at the N-terminus and to an ER-retention signal (HDEL) at the C-terminus, and the *OCS* terminator); *MYB98pro∷mRuby3–LTI6b* (the 1,610 bp *MYB98* promoter and *mRuby3* fused to the start codon of *LTI6b* (At3g05890) with the (SGGGG)_2_ linker, and the *HSP* terminator); *AKVpro∷H2B–mScarlet–I* (the 2,949 bp upstream regions of *AKV* (At4g05440; Boisnard-Lorig et al., 2001) and the full-length coding region of *H2B* (HTB1: At1g07790) fused to *mScarlet–I* with the (SGGGG)_2_ linker).

*SBT4.13pro∷SBT4.13–mClover* (coded as pDM349); The 2,040 bp upstream region and the full-length coding region of *SBT4.13* (At5g59120) were amplified and cloned into the pPZP221Clo using SmaI site (Takeuchi and Higashiyama, 2016).

*MYB98pro∷NLS–mRuby2* (coded as pDM371), a DNA fragment of NLS–mRuby2 (obtained from Addgene plasmid 40260), was amplified and then cloned into the pENTR/D-TOPO vector (Invitrogen, Japan), to generate pOR006. LR recombinations between the pDM286 (Maruyama et al., 2015) and pOR006 were performed using the LR clonaseII (Invitrogen) to produce pDM371.

The binary vectors were introduced into the *Agrobacterium tumefaciens* strain EHA105. The floral-dip or simplified *Agrobacterium*-mediated methods were used for the *Arabidopsis* transformations (Narusaka et al., 2010).

### 2.2 Microscopy

To image the female gametophyte development, we used two spinning-disk confocal microscope systems following the settings of Gooh et al. (2015), with the following modification: For the live imaging of the *in vitro* female gametophyte development, the confocal images were acquired using an inverted fluorescence microscope (IX-83; Olympus), equipped with an automatically programmable XY stage (BioPrecision2; Ludl Electronic Products Ltd, Hawthorne, NY, USA), a disk-scan confocal system (CSU-W1; Yokogawa Electric), 488-nm and 561-nm LD lasers (Sapphire; Coherent), and an EMCCD camera (iXon3 888; Andor Technologies, South Windsor, CT, USA). Time lapse images were acquired with a 60 silicone oil immersion objective lens (UPLSAPO60XS, WD = 0.30 mm, NA = 1.30; Olympus) mounted on a Piezo focus drive (P-721; Physik Instrumente). We used two band-pass filters, 520/35 nm for the GFP, and 593/46 nm for the tdTomato. The images were processed with Metamorph (Universal Imaging Corp.) and Fiji (Schindelin et al., 2012) to create maximum-intensity projection images and to add color.

We also used an inverted confocal microscope system with a stable incubation chamber (CV1000; Yokogawa Electric) equipped with 488 nm and 561 nm LD lasers (Yokogawa Electric), and an EMCCD camera (ImagEM 1K C9100-14 or ImagEM C9100-13; Hamamatsu Photonics, Shizuoka, Japan). Time lapse images were acquired with a 40*×* objective lens (UPLSAPO40*×*, WD = 0.18 mm, NA = 0.95; Olympus). We used the two band-pass filters, 520/35 nm for the GFP, and 617/73 nm for the tdTomato.

### 2.3 Isolation of female gametophyte cells

We used an inverted fluorescence microscope (IX-71; Olympus, Japan) equipped with a three-charge-coupled device (CCD) digital camera (C7780; Hamamatsu Photonics Ltd., Japan). Images were acquired using a 40 objective lens (LUCPlanFl 40, WD = 2.7–4 mm, NA = 0.60; Olympus). The unfertilized ovules of each cell marker line were treated with enzyme solution (1 % cellulase [Worthington, USA], 0.3 % macerozyme R-10 [Yakult, Japan], 0.05 % pectolyase [Kyowa Kasei, Japan], and 0.45 M mannitol [pH 7.0]). To collect the target cells, we used a micromanipulator (MN-4, MO-202U; Narishige, Japan) and micropipette (Picopipet HR; Nepa Gene, Japan) with glass capillaries (G-1; Narishige, Japan), which were pulled with a micropipette puller (P-97; Sutter, USA) (Ikeda et al., 2011).

### 2.4 cDNA preparation and library construction for sequencing

The mRNA was extracted from 12–18 synergid, egg, and central cells with Dynabeads mRNA DIRECT Micro Kit (Invitrogen, USA). Extracted mRNA were amplified using Ovation RNA-seq System V2 (NuGEN, USA). The RNA-seq libraries were prepared using a TruSeq RNA Sample Preparation Kit and Multiplexing Sample Preparation Oligonucleotide Kit (Illumina, USA). The libraries were sequenced on an Illumina GAIIx (Illumina) using 36 bp single-end reads.

### 2.5 RNA-seq data analysis

Reads were filtered by fastp (ver. 0.20.0; (Chen et al., 2018)). The cleaned reads were mapped to the Arabidopsis reference genome TAIR10, using HISAT2 (ver. 2.1.0; Kim et al. (2019)). The expression level for each gene was quantified as the read count and TPM with Stringtie (ver. 2.1.1; Pertea et al. (2015, 2016)). Differentially expressed genes between the synergid cells of the wild type and the *myb98* mutant were identified by TCC with a false discovery rate < 0.01 (ver. 1.24.0; Sun et al. (2013)). The TCC+baySeq (ver. 2.18.0) method with a false discovery rate < 0.01 was used for the identification of the differentially expressed genes among the synergid, egg, and central cells of the wild type (Osabe et al., 2019). Hierarchical clustering of the gene expression data was carried out using phylogram package (https://github.com/rambaut/figtree/).

## 3 RESULTS

### 3.1 Live imaging of the nuclear dynamics during female gametophyte development

The development of female angiosperm gametophytes *in vivo*, occurred within multiple layers of the maternal tissues of the flower. Christensen et al. (1997) defined the developmental stage by the observation of the fixed ovules (Figure 1A). Stage FG1 is the one-nucleate stage. Then the functional megaspore divided into two nuclei, without cytokinesis, during the first mitosis leading to the FG2 stage. A large vacuole then appeared at the center of the female gametophyte, separating the two nuclei to the micropylar and chalazal ends in the FG3 stage. After the second mitosis, the chalazal and micropylar nuclei migrated a line that is orthogonal to the chalazal–micropylar axis, at the early FG4 stage. The chalazal and micropylar nuclei migrated along a line that is parallel to the chalazal–micropylar axis at the late FG4 stage. After the third mitosis, the eight nuclei coenocyte is cellularized into the seven-celled female gametophyte in the FG5 stage. At the FG6 stage, the two polar nuclei fused to produce the secondary nucleus. Finally, the mature female gametophyte has two synergids, an egg cell, a central cell, and three antipodal cells. To investigate the actual developmental time course of the female gametophyte, we performed live-cell imaging of the female gametophytes development using the previously developed the *in vitro* ovule culture system for embryogenesis, using *Arabidopsis* (Gooh et al., 2015).

**Figure 1:**
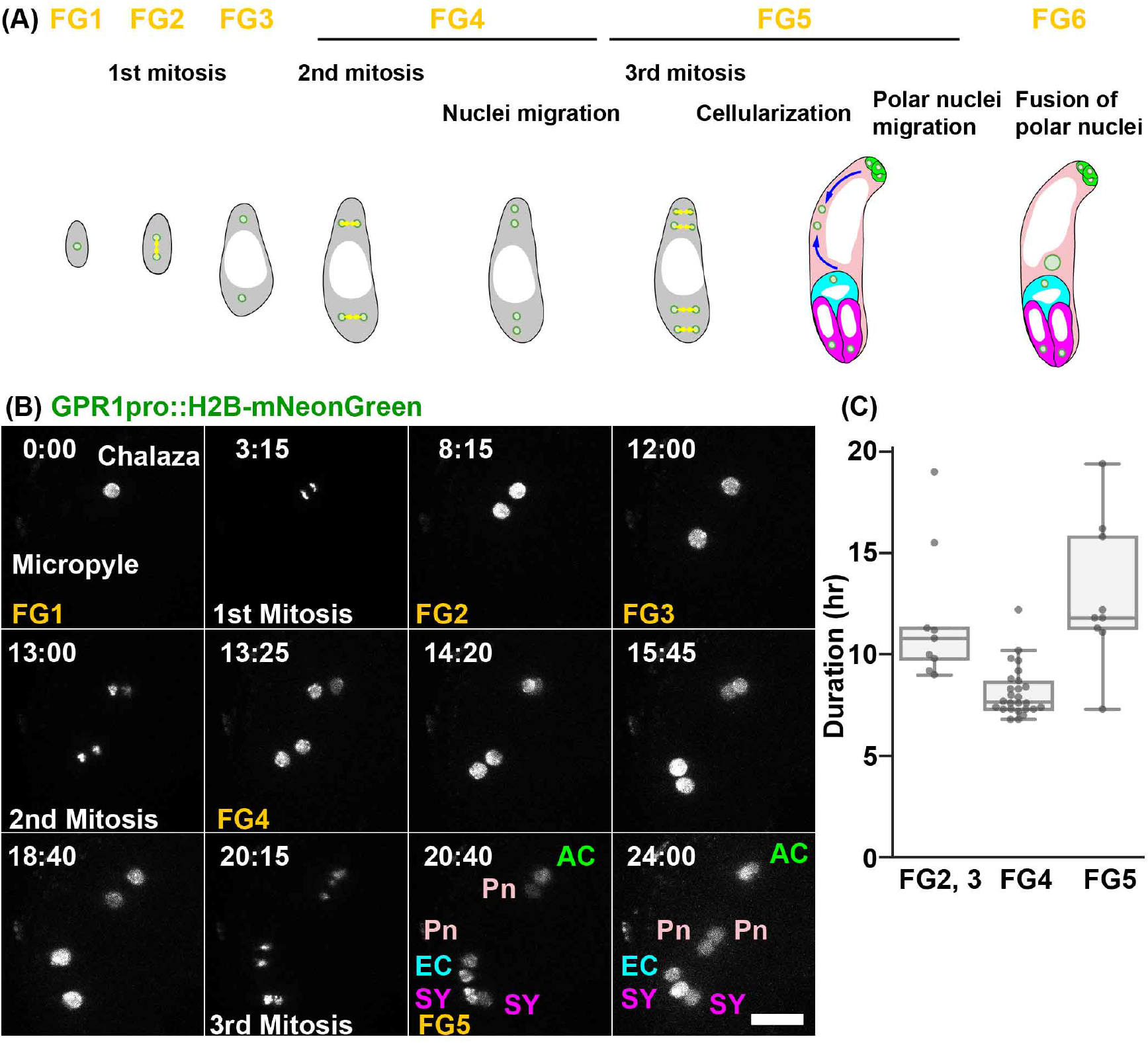
**(A)** Schematic representation of the development of a *Polygonum*-type female gametophyte. **(B)** Nuclei were labeled with *GPR1pro∷H2B–mNeonGreen*. The numbers indicate time (hr:min) from the onset of the observation. We succeeded in time-lapse recordings of the nuclear divisions in the isolated ovules from the FG1 to FG6. FG1, uninucleate functional megaspore; FG2, two-nucleate stage; FG3, two nuclei separated by a large central vacuole; FG4, four-nucleate stage; FG5, eight-nucleate/seven-celled stage; FG6, seven-celled with polar nuclei fused. AC, antipodal cells; EC, egg cell; Pn, Polar nucleus; SY, synergid cell. Scale bar, 20 *μ*m. **(B)** Durations of the nuclear divisions between the stages from FG2 to FG6. The interval times of the nuclear divisions for the female gametophyte development were analyzed for *GPR1pro∷H2B-mNeonGreen*, *RPS5Apro∷H2B-tdTomato*, and *RPS5Apro∷H2B-sGFP*.

To observe the nuclear dynamics in the female gametophytes development, we constructed GPR1pro∷H2B–mNeonGreen∷GPR1ter (Figure 1B, Supplementary Movie 1). GPR1 (GTP-BINDING PROTEIN RELATED1) was previously found to be expressed in the megaspore mother cells (i.e., at stage FG0) and the female gametophytes at FG1 −FG7 (Yang et al., 2017). At FG1, the nucleus was located at the center of the female gametophyte (Figure 1B; 0:00). Approximately 3 hr after the observation, the nucleus divided into two during the first mitosis (Figure 1B; 3:15). At FG2, the two nuclei were positioned at the center of the female gametophyte. Approximately 8 hr after the start of the FG2, each nucleus moved to the opposite ends of the ovule (Figure 1B; 12:00), at which point the vacuole may appear (FG3) (Christensen et al., 1997). After the second mitosis, the nuclei divided to lie in an orthogonal line along the chalazal–micropylar axis (Figure 1B; 13:00, 13:25). The chalazal nuclei migrated along a line that was parallel to the chalazal–micropylar axis (Figure 1B; 14:20), while the micropylar nuclei migrated along the surface of the female gametophyte, not parallel to the chalazal–micropylar axis(Figure 1B; 15:45, 18:40). The micropylar nuclei tended to lie along the abaxial surface of the female gametophytes (52/62, 84%). After the end of the third mitosis, the polar nuclei migrated linearly, not along the surface of the female gametophyte, towards each other to fuse (Figure 1B; 20:40, 24:00). We calculated the duration of each nuclear division from 30 movies of *GPR1pro∷H2B–mNeonGreen* (n = 19), *RPS5Apro∷H2B–tdTomato* (n = 10), and *RPS5Apro∷H2B–sGFP* (n = 1) (Figure 1C). The duration of the second and third nuclear divisions were 11.8 *±* 3.3 hr (mean *±* standard deviation; n = 9, Figure 1C; FG2,3) and 8.1 *±* 1.2 hr (n = 26, Figure 1C; FG4), respectively. After cellularization, it took 4.5 *±* 1.4 hr (n = 27) and 13.0 *±* 3.6 hr (n = 9, Figure 1C; FG5) after the third mitosis, for the polar nuclei to attach and fuse, respectively. Thus, the normal female gametophyte development was observed using the *in vitro* ovule culture system (Christensen et al., 1997).

### 3.2 Live imaging of the plasma membrane formation during female gametophyte development

To analyze the morphological changes in the female gametophytes, we observed their plasma membranes by labeling them with *RPS5Apro∷tdTomato–LTI6b* (Figure 2A, Supplementary Movie 2). The female gametophytes were located at the center of the ovule in the early stages of the female gametophyte development (Figure 2A; 17:20). The female gametophytes showed polar elongation towards the micropylar ends of the ovule (Figure 2A; 17:20, 11:00, 5:00). The fluorescent signals of the *RPS5Apro∷tdTomato–LTI6b* were detected in the plasma membranes of the female gametophytes during cellularization (Figure 2A; 0:00, arrow). Cellularization of the egg and synergid cells finished after 45 min and 1 hr 55 min, respectively (Figure 2A). The time differences between the cellularization of the egg and the synergid cells was 0.8 ± 0.2 hr (n = 10; Figure 2B). After the cellularization, the egg and synergid cells were elongated towards the chalazal end (Figure 2A; 3:50, 7:35). It took 4.0 ± 0.6 hr (n = 10) from the completion of the cellularization to the start of the elongation (Figure 2B).

**Figure 2:**
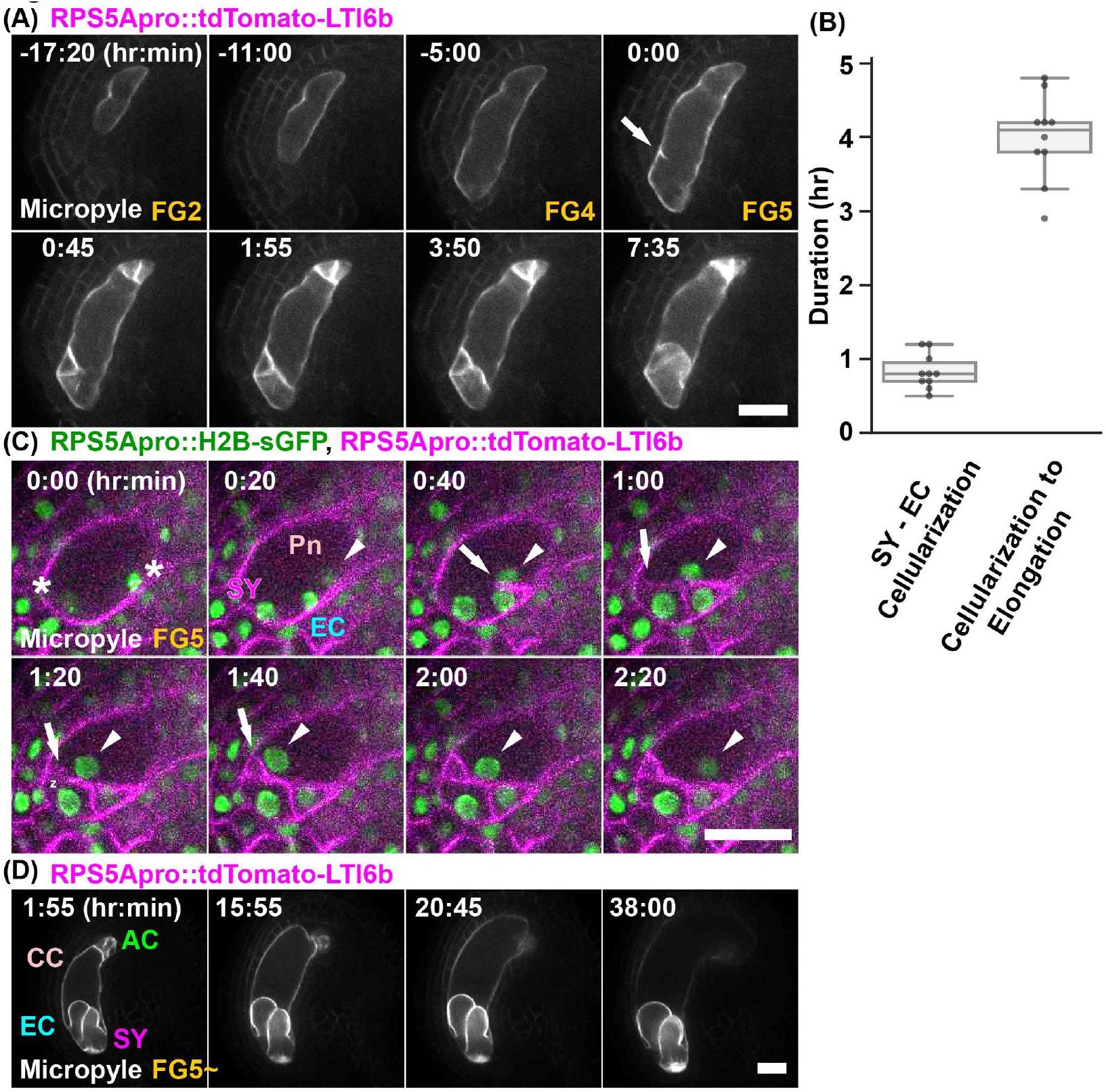
**(A)** Plasma membranes were labeled with *RPS5Apro∷tdTomato–LTI6b*. Numbers indicate time (hr:min) from the detection of the fluorescent signal of the tdTomato–LTI6b, on the forming cell plate (arrow). **(B)** Difference in the time to completion of the cellularization between the egg cell and synergid cells (left) and the initiation of the cell elongation from the completion of the cellularization (right) at the FG5 stage. **(C)** Nuclei and plasma membranes were labeled with *RPS5Apro∷H2B–sGFP* (green) and *RPS5Apro∷tdTomato–LTI6b* (magenta), respectively. Asterisks indicate the two micropylar nuclei at FG4. Arrows indicate the forming cell plate. Polar nucleus (arrowheads) migrated along the forming cell plate. **(D)** Plasma membranes were labeled with *RPS5Apro∷tdTomato–LTI6b*. Numbers indicate time (hr:min) from the onset of the observations. AC, antipodal cells; CC, central cell; EC, egg cell; Pn, Polar nucleus; SY, synergid cell. Scale bars, 20 *μ*m.

To analyze the relationship of the nuclear dynamics and the plasma membrane formation during the cellularization, we observed the *RPS5Apro∷tdTomato–LTI6b, RPS5Apro∷H2B–sGFP* ovule at the beginning of FG5 (Figure 2C, Supplementary Movie 3). In the case of the micropylar end, the fluorescent signals of the tdTomato–LTI6b were detected at the side nearest the nuclei, that gives rise to the polar nucleus and the egg nucleus after cellularization (Figure 2C; 0:40). This fluorescent signal was elongated to the opposite sides of the cell membranes of the female gametophytes. The polar nuclei migrated toward the opposite sides along with the plasma membrane formation (Figure 2C; 0:40 – 1:40). In the case of the chalazal end, the fluorescent signals of the tdTomato–LTI6b were also detected between the polar nucleus and the antipodal nucleus (Supplementary Movie 3, later). Thus, the dynamics of the plasma membrane formation were similar at the micropylar and chalazal ends.

During the maturation of the female gametophyte cells at the FG5 and FG6 stages, the central cell showed polar elongation towards the chalazal end of the ovule (Figure 2D, Supplementary Movie 4). A bright field movie showed that the central cell elongated by collapsing the chalazal regions of the ovule (Supplementary Movie 4). This direction of the elongation was the opposite to that of the FG2 – FG4 (Figure 2A; −17:20, −11:00, −5:00). As shown in Supplementary Movie 4, the antipodal cells appeared to be collapsing during the maturation of the central cell. However, we could not determine whether the antipodal cells degenerate or not, i.e., whether they reached FG7 (four-celled stage) or not (Song et al., 2014) in the *RPS5Apro∷tdTomato–LTI6b*. Although we could not observe the signature of FG7, such as degeneration of the antipodal cells, our *in vitro* culture system could monitor the entire development of the female gametophyte.

### 3.3 Live imaging of cell fate specification during female gametophyte development

The transcriptome data of the mature ovules indicated that each female gametophyte cell had specific gene expressions (Yu et al., 2005; Jones-Rhoades et al., 2007; Steffen et al., 2007). To investigate the initiation timing of the cell fate specification, we observed the mitochondria marker, the *EC1.2pro∷mtKaede* (Hamamura et al., 2011), in the ovules of the egg cells and the *MYB98pro∷GFP* (Kasahara et al., 2005) ovules of the synergid cells (Figure 3A,B, Supplementary Movie 5, 6). The fluorescent signals of the *EC1.2pro∷mtKaede* were detected in the egg cells before their elongation (Figure 3A; 0:00). Considering that the duration from egg cell cellularization to egg cell elongation was about 4 hr (Figure 2B), the *EC1.2* expression was initiated less than 4 hr after egg cell cellularization (Figure 2B). After 15.5 hrs, the fluorescent signals of the *ABI4pro∷H2B–tdTomato* were detected in the nucleus of the egg cell (Figure 3A; 21:10, arrowhead). Since MYB98 is an essential transcription factor for synergid cell function, the expression of MYB98 was predicted to begin after the synergid cells became cellularized; however, the fluorescent signals of the *MYB98pro∷GFP* were detected in the 4-nucleate female gametophytes at FG4, before the third mitosis and cellularization (Figure 3B; 3:10). After the cellularization, the fluorescent intensities of the GFP signals were increased in all of female gametophyte cells (Figure 3B; 0:40). As the cells mature, the GFP signals were decreased in the egg, central, and the antipodal cells, while they were increased in the synergid cells (Figure 3B; 8:20).

**Figure 3:**
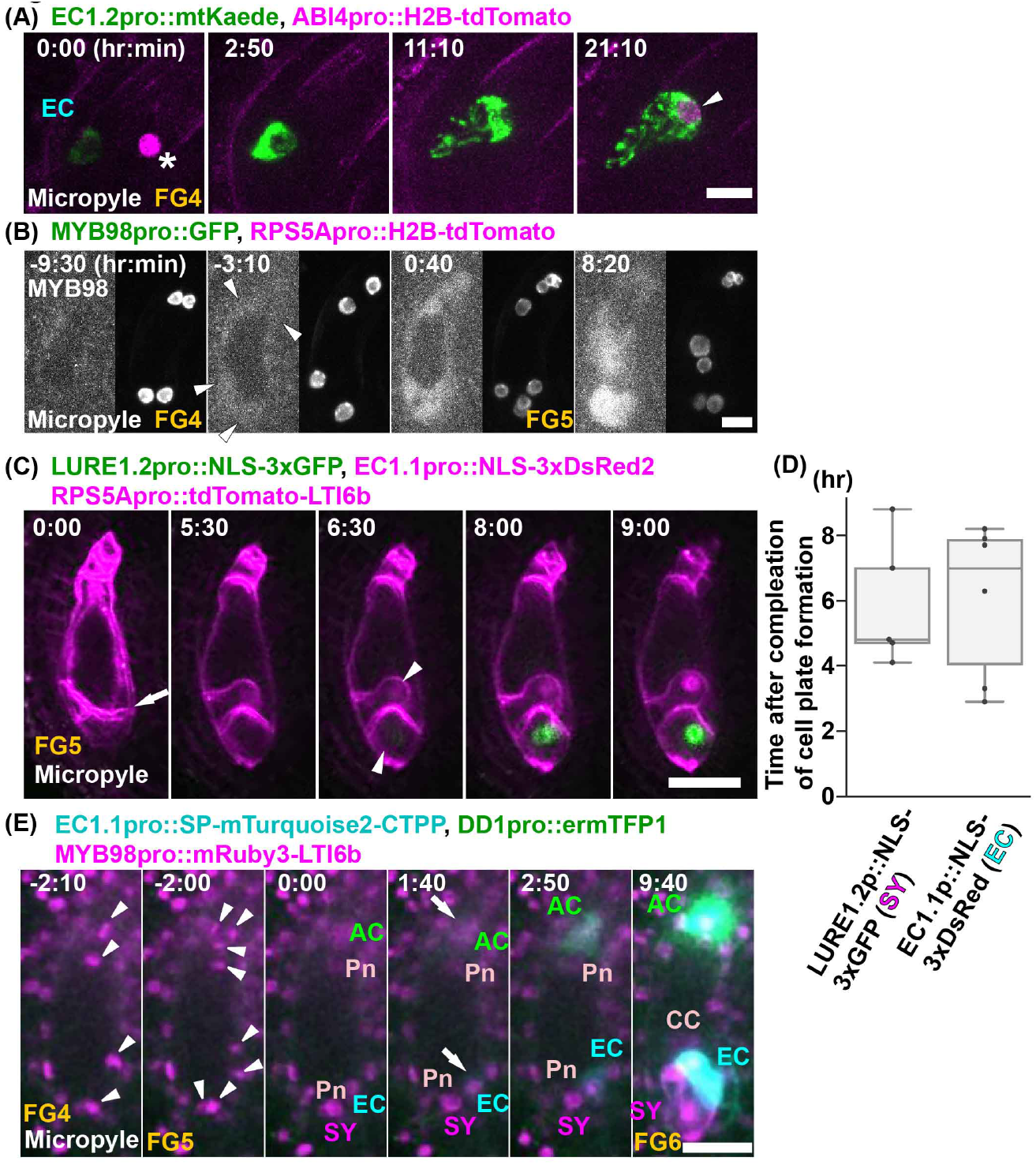
**(A)** The fluorescent signals of *EC1.2pro∷mtKaede* were observed for the egg cell fate. Nuclei were labeled with *ABI4pro∷H2B–tdTomato* (magenta). Numbers indicate time (hr:min) from the onset of observation. Asterisk indicates the background signal in the ovule (0:00). Arrowhead indicates the fluorescent signal of *ABI4pro∷H2B–tdTomato*. **(B)** The fluorescent signals of *MYB98pro∷GFP* were observed for the synergid cell fate. Nuclei were labeled with *RPS5Apro∷H2B–tdTomato*. Numbers indicate time (hr:min) after finishing the polar nuclear movement along the forming cell plate. Arrowheads indicate the first detection of the *MYB98pro∷GFP* signals. **(C)** Nuclei were labeled with *EC1.1pro∷NLS–3xDsRed2* (magenta) in the egg cells and *LURE1.2pro∷NLS–3xGFP* (green) in the synergid cells, respectively in the *FGR8.0*. The plasma membranes were labeled with the *RPS5Apro∷tdTomato–LTI6b* (magenta). Numbers indicate the time (hr:min) after finishing the cell plate formation. Arrow indicates the fluorescent signals of tdTomato–LTI6b on the forming cell plate. Arrowheads indicate the initiation of the expression of each cell-specific markers (6 hr 30 min). **(D)** Initiation of the expression of the cell-specific markers at FG5. The fluorescent signals for *EC1.1pro∷NLS–3xDsRed2* in the egg cells and *LURE1.2pro∷NLS–3xsGFP* in the synergid cells were observed after completion of the cell plate formation **(D)**. **(E)** Numbers indicate time (hr:min) from the third mitosis. Arrowheads indicate the chromosomes during the third mitosis. Arrows indicate the initiation of the expression of the specific markers of the egg cell (cyan) and the antipodal cells (green) 1 hr 40 min after cellularization. This timing was before the cell expansion and the polar nuclei migration. The *MYB98pro∷mRuby3–LTI6b* was detected 6 hr 20 min after cellularization. Scale bars, 10 *μ*m (A), 20 *μ*m (B).

To determine when the expression of each cell-specific marker began after cellularization, we utilized the female gametophyte-specific markers *FGR8.0* (Völz et al., 2013) and *RPS5Apro∷tdTomato–LTI6b* (Figure 3C, Supplementary Movie 7). After cellularization (Figure 3C; 0:00) and elongation of the egg and synergid cells (Figure 3C; 5:30), the *EC1.1pro∷NLS–3xDsRed2* and *LURE1.2pro∷NLS–3xGFP* signals were detected in the egg and synergid cells, respectively in *FGR8.0* (Figure 3C; 6:30). It took 5.9 ± 2.0 hr (n = 5) for the *EC1.1pro∷NLS–3xDsRed2* to be detected after the completion of the cellularization (Figure 3D). Considering that the expression of the *EC1.2pro∷mtKaede* was initiated before the egg cell elongation (Figure 3A), the detection of the NLS marker was slower than that of the mitochondrial marker.

To investigate the correlation between the timing of the expressions of each cell-specific markers at the FG5, we used the multiple cell-type-specific marker line (Figure 3E, Supplementary Movie 8). We changed the target signals of the new markers from the NLS and the fluorescent proteins as detection may have been slow. The cell-specific markers of the egg cell (*EC1.1pro∷SP–mTurquoise2–CTPP*) and the antipodal cells (*DD1pro∷ermTFP1*) were expressed 1.7 hr after cellularization (Figure 3E; 1:40). This was before the egg and synergid cell elongations and the polar nuclei migrations. These results suggested that each cell fate was specified almost immediately after cellularization at the eight-nucleate stage.

### 3.4 *myb98* synergid cells showed aberrant morphology and subcellular dynamics

MYB98 is required for the formation of the filiform apparatus during the synergid cell differentiation and the expression of the AtLURE1 peptides to attract the pollen tube (Kasahara et al., 2005; Takeuchi and Higashiyama, 2012). However, *MYB98pro∷GFP* was detected before cellularization in FG4 and all of the female gametophyte cells in FG5 (Figure 3B). To clarify the effects of the MYB98 transcription factor on the female gametophyte specifications, we observed the morphology and nuclear dynamics with the promoter activity of the *MYB98* in the synergid cells of the wild type and *myb98* mutant ovules (Figure 4, Supplementary Movie 9, 10). The fluorescent signals of the *MYB98pro∷NLS–mRuby2* were also detected in all of the female gametophyte cells, as well as the synergid cells of the wild type and *myb98* ovules. However, the *MYB98pro∷NLS–mRuby2* signals were detected during the synergid cell elongation, later than the *MYB98pro∷GFP*. These results indicated that the expression of NLS–mRuby2 was slower than that of the free GFP. Detection of *RPS5Apro∷tdTomato–LTI6b* in the forming cell plate to the egg cell elongation took 4.7 ± 0.6 hrs (Figure 2A,B). Considering that the mRuby2 and EGFP required 150 min and 25 min to mature, respectively (Lam et al., 2012), the NLS may take a long time to localize to the nucleus after transcription. Furthermore, since the NLS line had a considerable variations in the time required for the detection of the expression (Figure 3D), free-fluorescent proteins or other signal peptide-fusions were preferred to determine the timing required for the transcription. Although the nuclei were always located at the micropylar end of the synergid cells in the wild type (Figure 4A), it moved around in the synergid cells of the *myb98* (Figure 4B). The nuclei tracking over 14 hr also showed that the nuclei of the *myb98* moved closer to the chalazal end than to the wild type (Figure 4C). The large vacuoles occupied the chalazal end of the synergid cells in the wild type (Figure 4A). This polar distribution of the vacuole was disturbed in the synergid cells of the *myb98* (Figure 4B). In addition, the *myb98* synergid cells were more elongated during the maturation (Figure 4B; 2:50—8:20). The results showed that the absence of the *MYB98* affected the morphology and cellular dynamics of the synergid cells in addition to the formation of the filiform apparatus (Kasahara et al., 2005).

**Figure 4:**
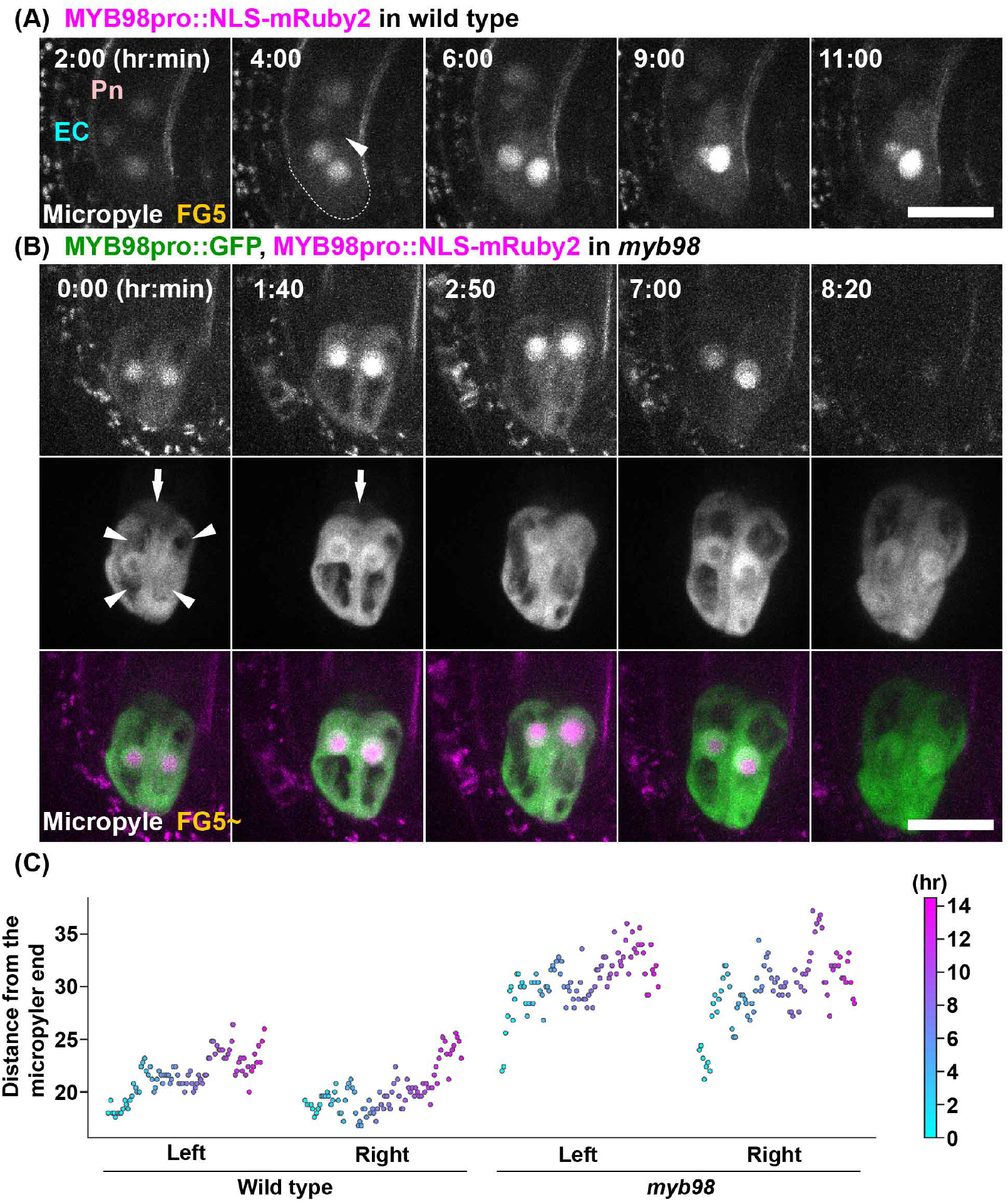
**(A)** Nuclei of the synergid cells were labeled with *MYB98pro∷NLS–mRuby2* in the wild type. The numbers indicate the time (hr:min) from the onset of the observations. Dashed lines indicate the surface of the synergid cells at the micropylar end. **(B)** Nuclei of the synergid cells that were labeled with *MYB98pro∷NLS–mRuby2* in the *myb98*. The fluorescent signals of the *MYB98pro∷GFP* were observed for the synergid cell fate. The arrowheads indicate the vacuoles in the synergid cells. The arrows indicate the GFP signals in the egg cells. Scale bars, 20 *μ*m. **(C)** Nuclei positions on the micropylar–chalazal axis were plotted in each synergid cell in the wild type and *myb98* from the Supplemental Movies S9 and S10. Each point color indicates the time corresponding to the color bar. The leftmost point indicates the start time. The y-axis indicates the distance from the micropylar end of the synergid cell.

### 3.5 Gene expression analysis of the female gametophyte cell

To investigate the gene expression profiles of the synergid cells in the wild type and *myb98* mutant, we established a method to isolate them in *Arabidopsis*. We treated the ovules in emasculated ovaries of the transgenic marker line for the synergid cells, *MYB98pro∷GFP* (Figure 5B), with enzyme solutions. The protoplasts of the synergid cells were released from the ovules through their micropyles with enzyme treatment for 30–60 min (Figure S1A), and those with the GFP signals were collected by micromanipulation (Figure 5A). Initially, the synergid cell-derived protoplasts mostly associated with other GFP-negative ovular cells, probably due to insufficient cell wall digestion. To increase the efficiency of the single synergid cell isolation, we optimized the following two conditions. One was the calcium nitrate in the enzyme solution as the calcium ion was suggested to inhibit the degradation of the cell wall (Imre and Kristóf, 1999). Subsequently, the removal of calcium ion from the enzyme solution decreased the adhesion of protoplasts and increased the frequency of the collectable synergid cells that were released as single cells (Figure S1B, Table S1). The other condition was the pH of the enzyme solution. We found that the protoplasts began to decrease the GFP fluorescence in a short period and eventually ruptured after the cell surface that gradually became rough, and this may be related to the decreases in viability. We performed the enzyme treatments at pH values of 5.0–9.0 and observed the GFP fluorescence as a vital indicator of the protoplast (Chiu et al., 1996). The rate of the GFP-positive synergid protoplasts was highest at pH 7.0, which was the best for the isolation of the synergid cells (Figure S1C, Table S2). The optimized enzyme solutions allowed us to collect pure synergid cells with high efficiency (Figure 5C). To isolate other types of female gametophyte cells, we examined the enzyme solution treatment with the ovules of each marker line, *EC1.2pro∷mtKaede* and *FWApro∷FWA–GFP*, for the egg and central cells, respectively (Hamamura et al., 2011; Kinoshita et al., 2004). The protoplasts of the two gametic cells were also detached from their ovules through the micropyle (Figure 5E–H).

**Figure 5:**
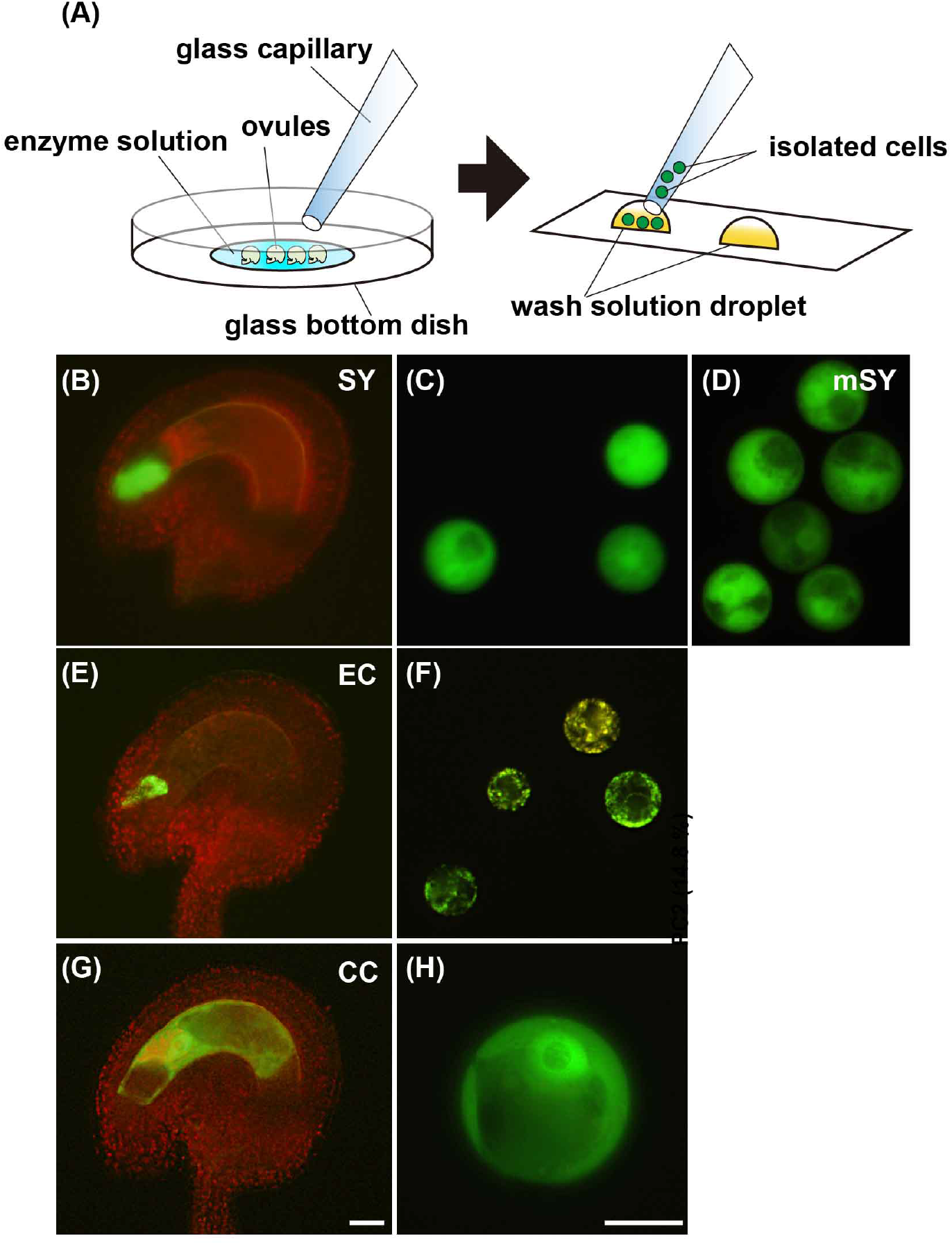
Isolation of the female gametophyte cells. **(A)** Scheme for the isolation of the female gametophyte cells. The ovules of the marker lines for the synergid **(B)**, egg **(E)**, and central cells **(G)**. The isolated each type of cell. Synergid cells in the wild type **(C)** and *myb98* mutant **(D)**. **(F)** Egg cells. **(H)** Central cell. SY, synergid cell; EC, egg cell; CC, central cell; mSY, synergid cell of *myb98* mutant. Scale bars, 20 *μ*m.

We then performed RNA-seq to analyze the gene expression profiles of the collected the synergid, egg, and central cells in the wild type and the synergid cells in the *myb98* mutant (Figure 5D). RNA-seq data from these female gametophyte cells were mapped to the genome of *Arabidopsis* (TAIR version 10) with the published sequence data from the ovules at 12 hr-after-emasculation (HAE) (Kasahara et al., 2016) and 2-week-old seedlings (Rogers et al., 2012). There were 4,996–18,432 genes (read counts > 10) detected in each sample (Figure 6A; Table S3). Hierarchical clustering showed that all samples were clustered into six independent groups (Figure 6B). The principal component analysis (PCA) indicated that the PC1 (34.2 %) and PC2 (14.8 %) were sufficient for separating these samples into the six groups (Figure 6C). These results suggested that our datasets had a high level of reproducibility. The expression profile of the synergid cells in the mutant was more like that of the egg cells than the synergid cells in the wild type. We identified the differentially expressed genes (DEGs) among the central cell, the egg cell, and the synergid cells in the wild type and between the synergid cells in the wild type and *myb98* mutant (Table S3). Interestingly, several egg cell-specific genes were highly expressed in the mutant synergid. We examined the expression patterns of the DEGs in the synergid dataset among all samples (Figure 6D). The cluster of mutant synergids was closer to that of the egg cells than the synergid cells in the wild type. These results also indicated that the expression pattern of the *myb98* mutant synergid was partially changed to be egg cell-like.

**Figure 6:**
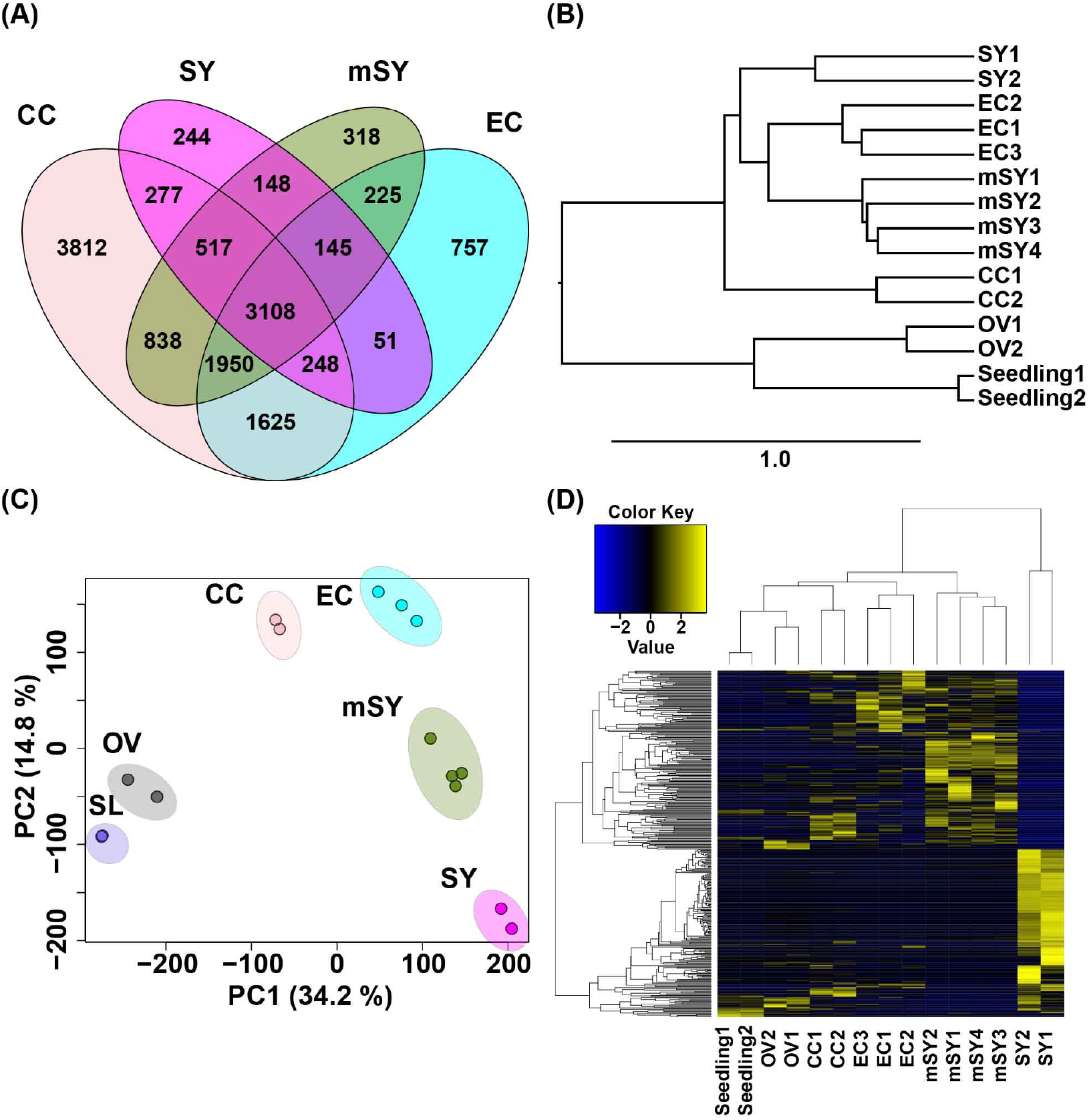
RNA-seq of the female gametophyte cells. The biological replicates were sequenced for the two SY, two CC, three EC, and four mSY cells. **(A)** Hierarchical clustering of samples for the RNA-seq. **(B)** The PCA analysis of all transcriptome data, the female gametophyte cells, ovules and seedlings. **C** Venn diagram of the expressed genes (> 10 reads) in each cell type. **D** Heatmap of the differentially expressed genes between the synergid cells in the wild type and the *myb98* mutant. OV, ovule; SL, seedling.

### 3.6 Egg cell-specific markers were expressed in one of the synergid cells of *myb98*

To confirm the expression patterns of the egg cellspecific genes in the *myb98*, we analyzed the CDR1–LIKE aspartyl proteases, which are highly expressed in the egg cells (Table S4). CDR1 (CONSTITUTIVE DISEASE RESISTANCE 1) was previously found to be involved in the peptide signaling of disease resistance (Xia et al., 2004). The phylogenetic analysis showed that *Arabidopsis* contained two distinct groups of CDR1s: a CDR1–LIKE2 (At1g31450)/CDR1–LIKE1 group (At2g35615) and a CDR1 (At5g33340)/CDR1–LIKE3 (At1g64830) group ((Olivares et al., 2011); Figure 7A). The *CDR1–LIKE2pro∷CDR1–LIKE2–mClover* (hereafter *CDR1L2–mClover*)and *CDR1–LIKE1pro∷CDR1–LIKE1–mClover* were expressed only in the egg cells, while the *CDR1pro∷CDR1–mClover* was expressed in the central and the antipodal cells (Figure 7B, Supplementary Movie 11). These localizations were consistent with the groupings of the CDR1s by the phylogenetic analysis. Although the fluorescent signals of *CDR1L2–mClover* were limited to the egg cell after cellularization in the wild type (Figure 7B, Supplementary Movie 12, Table 1; 100%, *n* = 6), *myb98* mutant had supernumerary cells with CDR1L2–mClover signals at the micropylar end (Figure 7B,C, Supplementary Movie 12, Table 1; 100%, *n* = 9). Initially, the CDR1L2–mClover signal was limited to a single cell at the egg cell position (Figure 7C; 0:00). However, 9.5 hr after the signal detection in the egg cell, the *CDR1L2–mClover* signal was also detected in one of the synergid cells (Figure 7C; 9:30). In most cases, one of the synergid cells had the expression of *CDR1L2–mClover* in the *myb98* (Table 2; 89%, *n* = 9).

**Figure 7:**
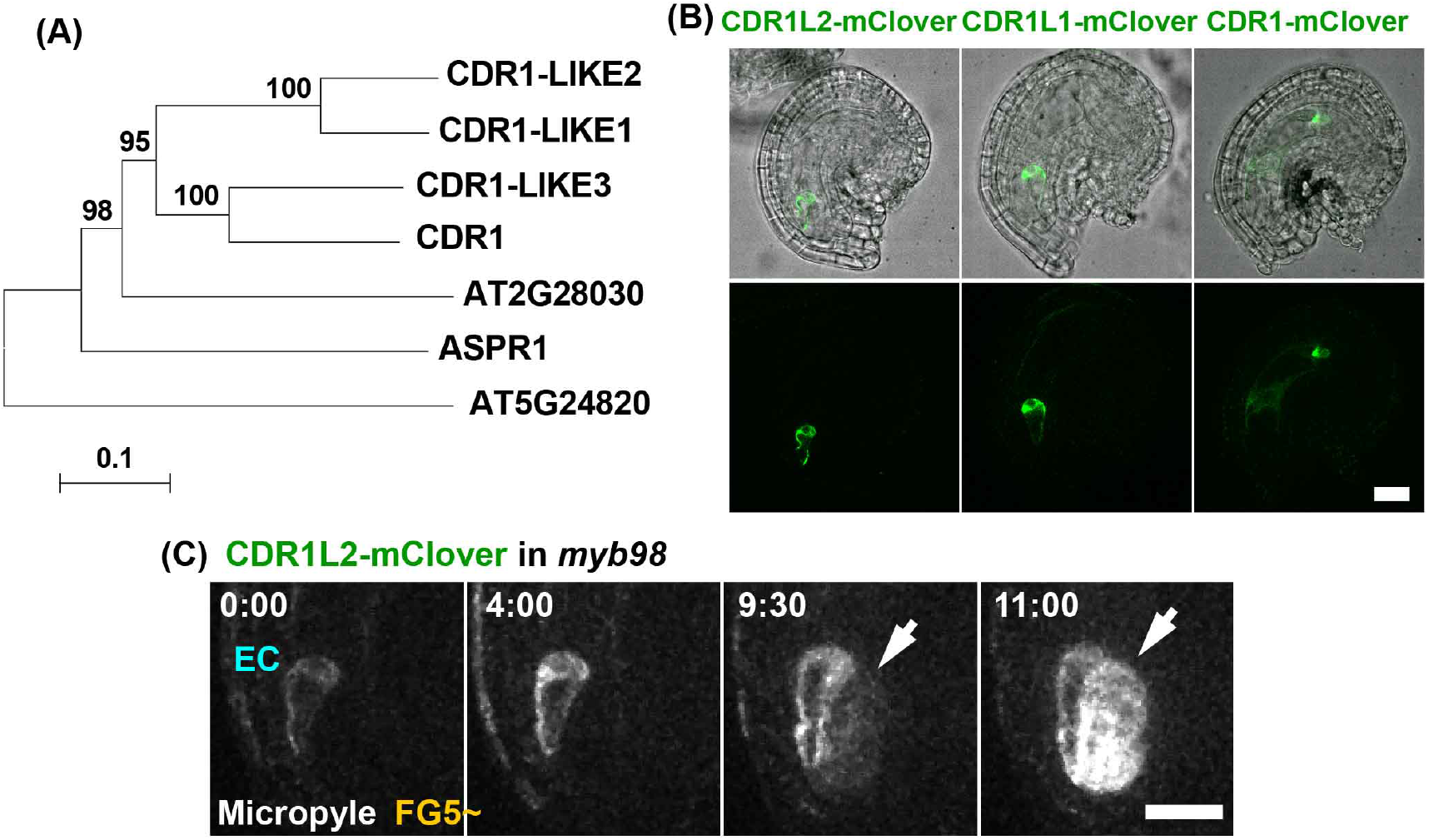
**(A)** Phylogenetic tree of the aspartyl proteases in the *Arabidopsis thaliana*. **(B)** The expression patterns of the *CDR1L2–mClover* and *CDR1L1–mClover* were detected in the egg cell. The fluorescent signal of the *CDR1–mClover* was detected in the central cell and the antipodal cells. **(C)** The expression of the *CDR1L2–mClover* in the *myb98* mutant ovules. The numbers indicate the time (hr:min) from the onset of the observations. The arrow indicates the CDR1L2–mClover signals in the synergid cell of the *myb98*. Scale bars, 20 *μ*m.

**Table 1:** Expression of EC-specific genes in the female gametophyte cells

| Construct | MYB98<br>genotype | Expression |  |  |
| --- | --- | --- | --- | --- |
|  |  | EC | EC, SY | EC, SY, AC |
| CDR1-LIKE2pro:: | +/+ | 6/6 (100 %) | 0 | 0 |
| CDR1-LIKE2-mClover | -/- | 0 | 9/9 (100%) | 0 |
| SBT4.13pro:: | +/+ | 10/10 (100%) | 0 | 0 |
| SBT4.13-mClover | -/- | 1/23 (4%) | 15/23 (65%) | 7/23 (30%) |

**Table 2:** Number of mutated synergid cells with EC-specific gene expressions

| Construct | MYB98<br>genotype | Expression |  |  |
| --- | --- | --- | --- | --- |
|  |  | EC | EC, SY | EC, SY, AC |
| CDR1-LIKE2pro:: | +/+ | 6/6 (100 %) | 0 | 0 |
| CDR1-LIKE2-mClover | -/- | 0 | 8/9 (89%) | 1/9 (1%) |
| SBT4.13pro:: | +/+ | 10/10 (100%) | 0 | 0 |
| SBT4.13-mClover | -/- | 1/23 (4%) | 17/23 (74%) | 5/23 (22%) |

Previously, the *myb98* mutant synergid cells were found to have high expression levels for the egg cell-specific gene, SBT4.13 (Bleckmann and Dresselhaus, 2016). The fluorescent signal of *SBT4.13pro∷SBT4.13–mClover* was detected only in the egg cell before the egg cell elongation (Figure 8A; 0:00–1:30, Supplementary Movie 13). This expression timing of the *SBT4.13pro∷SBT4.13–mClover* was similar to that of *EC1.2pro∷mtKaede* (Figure 3A). The *myb98* ovules showed two patterns of *SBT4.13pro∷SBT4.13–mClover* in the female gametophyte (Figure 8B, Supplementary Movie 14). One is the expression of *SBT4.13pro∷SBT4.13–mClover* in the synergid and the antipodal cells in addition to the egg cells of the *myb98* ovules (Figure 8B; upper, Supplementary Movie 14; former, Table 1; 30%). The other is the synergid and the egg cell (Figure 8B; lower, Supplementary Movie 14, later, Table 1; 65%). Similar to the results for the *CDR1L2–mClover*, one of the synergid cells showed *SBT4.13pro∷SBT4.13–mClover* expression in the *myb98* (Table 2; 74%).

**Figure 8:**
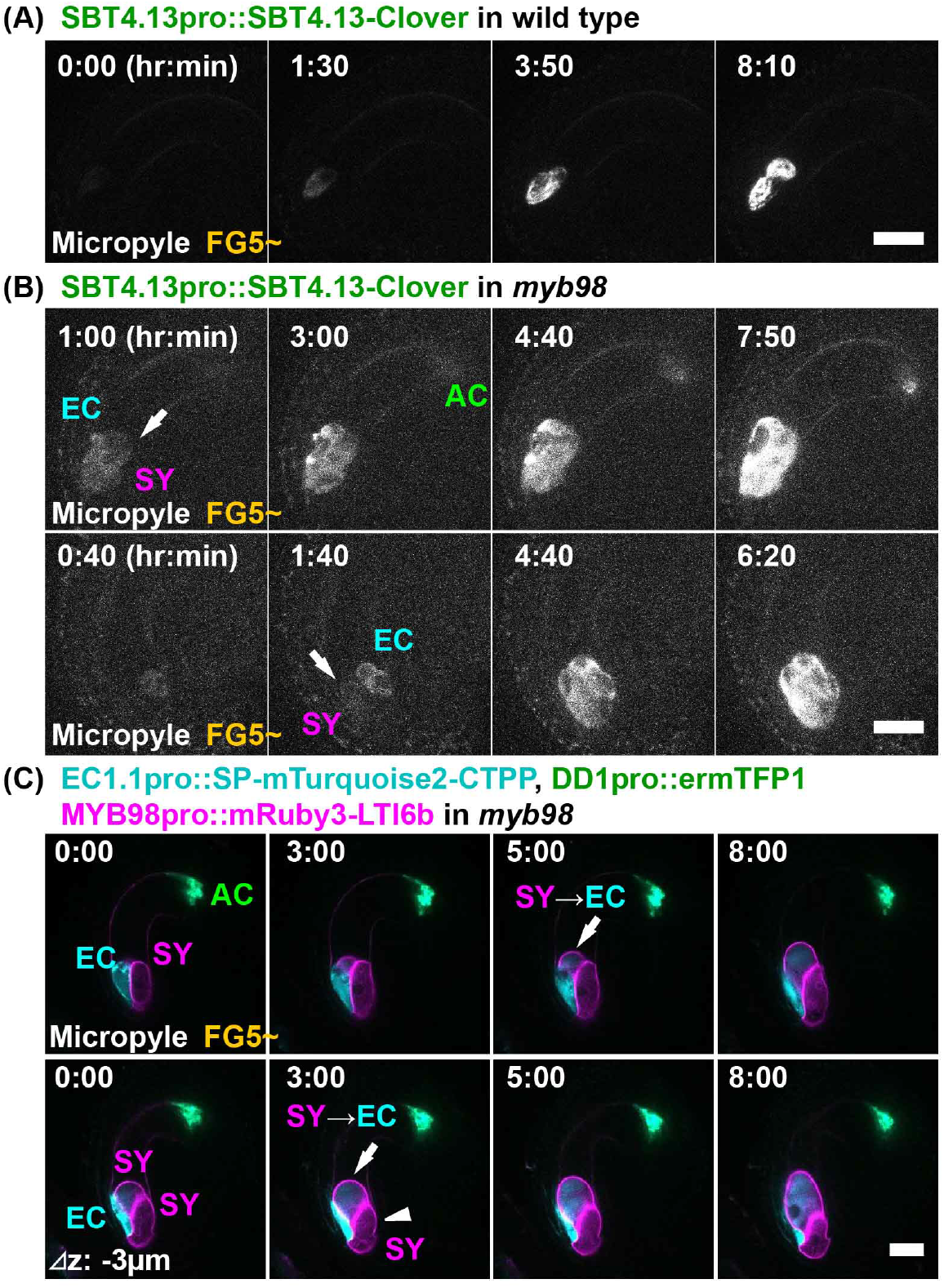
**(A,B)** The expression patterns of the *SBT4.13pro∷SBT4.13–mClover* in the wild type (A) and *myb98* (B) mutant ovules. The numbers indicate the time (hr:min) from the first detection of the SBT4.13–mClover. The fluorescent signals of the SBT4.13–mClover were only detected in the egg cells of the wild type. (A). However, in the case of the *myb98*, the fluorescent signals of the SBT4.13–mClover were also detected in the synergid cells (B; upper) and the antipodal cells (B; lower). **(C)** The expression patterns of the female gametophyte-specific markers in the *myb98*. The numbers indicate the time (hr:min) from the onset of the observations. At first, the *MYB98pro∷mRuby3–LTI6b* were detected in the two synergid cells (0:00). The arrows indicate the *EC1.1pro∷SP–mTurquoise2–CTPP* expression in one of the synergid cells (3 hr 00 min, 5 hr 00 min). The arrowhead indicate no expression of the *EC1.1pro∷SP–mTurquoise2–CTPP* (3 hr 00 min). The upper and lower panels are different *z* planes. Scale bars, 20 *μ*m.

To determine whether the egg cell-specific markers were expressed in the one or two synergid cells more clearly, we observed the *myb98* ovules in the multiple cell-type-specific marker line (Figure 8C, Supplementary Movie 15). After detection of the *MYB98pro∷mRuby3–LTI6b* signal in the two synergid cells (Figure 8C; 0:00), the signal of the *EC1.1pro∷SP–mTurquoise2–CTPP* was detected in one of the synergid cells (Figure 8C; 0:00; lower). Thus, one of the synergid cells showed that cell fate conversion to the egg cell in the *myb98*.

## 4 DISCUSSION

We established a live-female gametophyte imaging system to visualize the nuclear divisions and cell fate specifications in *Arabidopsis thaliana*. This system revealed the living-dynamics of the female gametophyte development, such as nuclear movements, cell elongation, duration of each FG stage, and the expression time of each cell-specific gene (Figure 9). Previously, we had developed the N5T medium for *in vitro* ovule cultures to perform live-cell analysis of the embryo development in *Arabidopsis* (Gooh et al., 2015). The Nitsch medium supplemented with 5% trehalose, resulted in the highest percentage of ovule survival *in vitro* during seed development after fertilization. This medium also enabled us to perform live-cell imaging during female gametophyte development, prior to fertilization. There are different technical advantages and limitations of the live-cell imaging of the female gametophyte development within the ovules in *Arabidopsis*.

**Figure 9:**
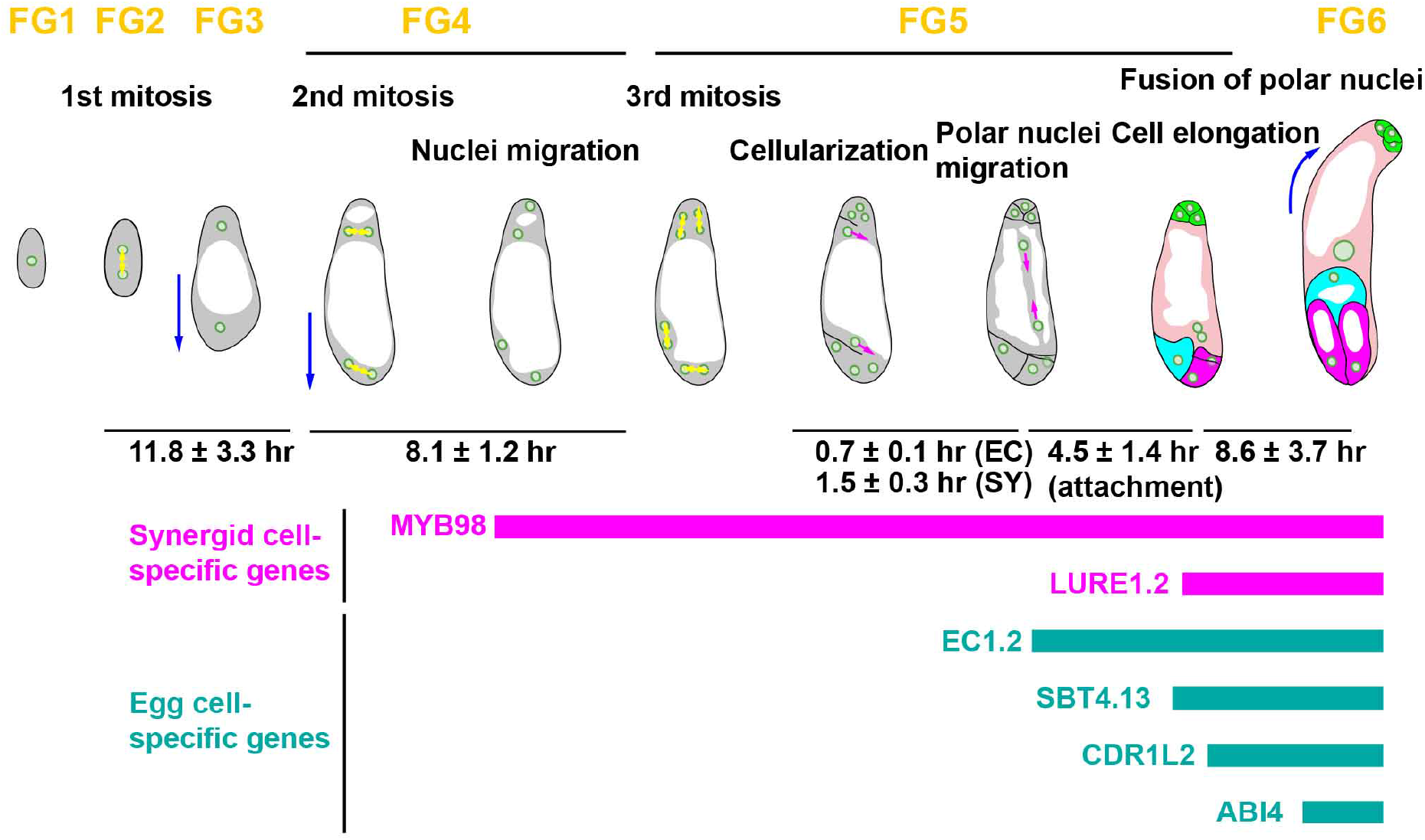
Schematic illustration of dynamics of the female gametophyte development in *Arabidopsis*. Yellow arrows show the direction of nuclear divisions. Blue arrows show the direction of cell elongation of the female gametophyte. Magenta arrows show polar nuclear migration at FG5. The time (mean ± standard deviation) calculated from the movies.

One advantage is the observation distance of the female gametophyte within the ovule. In the case of the embryo development, the ovule expansion during the seed development makes it difficult to observe the subcellular structures of the embryo within the ovule by confocal microscopy. Therefore, two-photon microscopy helps us to perform deep imaging of the zygote and embryo within the ovules. (Gooh et al., 2015; Kimata et al., 2016, 2019). The ovule elongated only along the micropylar–chalazal axis via the growth of the female gametophyte and the integument (Figure 2A,D). Therefore, it was possible to conduct live-cell imaging of the female gametophytes development with high resolution using confocal microscopy.

One limitation was the difficulties with the expansion of the female gametophytes during early development, as during *in vitro* ovule cultures, the female gametophytes collapsed in some cases. A possible cause was the change of turgor pressures in the female gametophytes. Optimal osmotic conditions for the isolation of the female gametophytic cells in *Torenia fournieri*, showed that the osmotic pressures increased from FG0 to FG4 and decreased from FG4 to FG6, at their peaks (Imre and Kristóf, 1999). The *T. fournieri* was slightly different from *Arabidopsis*, as the female gametophyte was naked from FG4, but it was inferred that the osmotic pressure was different during the female gametophyte development, even in the *Arabidopsis*. Especially in the early stages (FG0), the female gametophytes were not enclosed by the integuments. As a result, the *in vitro* developments of the integuments did not proceed, and the development of the female gametophytes was stopped. When the integuments covered the female gametophytes in the late FG0, the female gametophyte development proceeded *in vitro* (Figure 1B). To observe meiosis, megasporogenesis, and other early processes *in vitro* in real-time, it was considered that further improvements were required, such as the determination of conditions in which a placenta was attached without isolation.

### 4.1 Subcellular dynamics in female gametophyte development

To date, the female gametophyte of *Arabidopsis* has been analyzed only in fixed samples, so the actual developmental time course and subcellular dynamics were not known (Christensen et al., 1997). One of the major events that could not be seen in the fixed samples was that the vacuoles were dynamic in the female gametophytes. In the previous schematics, the vacuoles were drawn as large and only in the center of the cell (Drews and Koltunow, 2011). When the polar nuclei migrated to fuse to each other at FG5, they were described as moving along the periphery of the female gametophyte to avoid the large vacuole. (Sprunck and Groß-Hardt, 2011). However, the observations of the present study showed that the polar nuclei migrated linearly to fuse and adhere to the vacuole in the middle of the cell at shorter distances (Figure 1). This result suggests that the vacuoles of the female gametophyte did not remain large and static, but changed shape dynamically. The dynamics of the vacuoles have been seen in *Arabidopsis* and tobacco BY-2 cultured cells, and this plasticity is due to actin filaments (Higaki et al., 2006; Segami et al., 2014). As actin filaments were also involved in the nuclear migrations during gamete fusion, the linear migration of the polar nuclei was expected to involve actin filaments (Kawashima et al., 2014). In the mature central cells after polar nuclei fusions, the nucleus of the central cells were located to the micropylar end, and the actin filaments played an important role in the positioning of the nucleus (Kawashima and Berger, 2015). The vacuoles were located at the chalazal end of the synergid cells and the micropylar end of the egg cells, thus appearing to limit the nuclear migration (Figs.3A, 4A). In the case of *myb98* mutant, the vacuoles were dynamic, causing the nuclei to move around and not to stay in one place (Figure 4B,C). It is considered that this nuclear movement promoted the expression of the egg cell markers in the synergid cells of *myb98*. Alternatively, this movement may appear as a mixture of the egg and synergid cells identity (Figure 6C). Strong correlations between the nuclear position and the cell fate were shown in several mutants (Kong et al., 2015; Groß-Hardt et al., 2007; Pagnussat et al., 2007; Moll et al., 2008; Kirioukhova et al., 2011). However, it remains unclear whether the nuclear position determines gene expression or gene expression determines the nuclear positioning. Manipulation of nuclear behavior with the *in vitro* ovule culture systems will help to reveal the mechanisms of cell fate specifications in the development of the female gametophytes.

### 4.2 The synergid cells of *myb98* showed egg-cell like gene expressions

Previous studies have supported the lateral inhibition model for the differentiation of the female gametophyte cells. Although all cells in the female gametophyte have the gametic cell competence, the accessory cells like the synergid and antipodal cells, are repressed in the gametic cell fate (Groß-Hardt et al., 2007; Tekleyohans et al., 2017). In the present study, the RNA-seq of the female gametophyte cells identified many of the DEGs and the highly expressed genes in each type of cell (Table S3). We compared the DEGs between wild type and *myb98* identified by this RNA-seq study with those identified by the microarrays (Jones-Rhoades et al., 2007). The number of upregulated genes in the *myb98* was 204 and 40 from the RNA-seq and microarray, respectively (Figure S1D). The number of downregulated genes in the *myb98* was 188 and 77 from the RNA-seq and microarray, respectively (Figure S1E). These results suggested that cell-specific RNA-seq had much higher sensitivity for the detection the DEGs than the microarrays, because of the number of DEGs. Although 70 downregularted genes in *myb98* were overlapped between RNA-seq and microarray data, only 4 upregulated genes in *myb98* were overlapped (Figure S1D,E). The differences in the upregulated genes of the *myb98* may be caused by the wild type background or the developmental stage for the sampling (Jones-Rhoades et al., 2007). Furthermore, our RNA-seq revealed that the gene expression profiles of the *myb98* mutant synergid, changed partially to the egg cell-like (Table S4). A previous microarray analysis of *myb98* presented different results, however, this is thought to be because their sample contained the entire ovule, not the synergid cells alone (Jones-Rhoades et al., 2007). The RNA-seq conducted here allowed for the isolation of single cell types and mutants, and thus enabled the detection of cell-specific changes. This has evidenced the power of this method for investigation of cell fate specification mechanisms.

The *MYB98* was reported as the gene that controlled the characteristic development of the synergid cells (Kasahara et al., 2005). The *myb98* synergid was like a deficient egg cell, because an important factor for the synergid cell fate was lost. The hierarchical clustering and the difference of gene expressions reflect the intermediary state of the *myb98* synergid (Figure 6B,C,D). Further research is required to identify if the synergid cells of the *myb98* function as egg cells, synergid cells, or both.

### 4.3 The complex regulation is necessary for the egg cell specification and function

As the fluorescence of the *EC1.2pro∷mtKaede* was detected before the egg cell elongation, it was considered that the expression of the *EC1.2* began immediately after cellularization. *CDR1L2–mClover* and *ABI4pro∷ H2B–tdTomato* were expressed at the stage of egg cell maturation, whereas the *EC1.2pro∷mtKaede*, *EC1.1pro∷NLS–3xDsRed2*, and *SBT4.13pro∷SBT4.13–mClover* began to be expressed immediately after cellularization, and before egg cell elongation. The fact that egg cell-specific genes were expressed at different times provides clues as to their function and the regulation of their expression. The synergid cells of the *myb98* mutant also showed these differences in the timing of the expression in the egg cells. In the *myb98*, *SBT4.13pro∷SBT4.13–mClover* was expressed almost simultaneously in the egg cells, and the synergid cells during the cell elongation. On the other hand, *CDR1L2–mClover* was expressed in the synergid cells after egg cell maturation. The expression of *SBT4.13pro∷SBT4.13–mClover* in the synergid cells from the early stage indicated that the *myb98* synergid cells had changed their cell fate from the early stage. These results suggested that what each gene senses and recognizes as an egg cell is different. The *myb98* pistils had only one embryo after fertilization (10 pistils; 63 ovules). This result indicated that the synergid cells with the egg cell-specific genes were not functional for fertilization in the *myb98*. The additional egg-like cells appear to not be functional in the *lis*, *clo*, *ato* and *wyr*(Groß-Hardt et al., 2007; Moll et al., 2008; Kirioukhova et al., 2011). However, the *amp1* has twin embryos and *eostre* has twin zygote-like cells, indicating that these additional egg-like cells are functional for fertilization (Pagnussat et al., 2007; Kong et al., 2015). These differences in the gene expressions of the mutants may provide clues as to the acquisitions of the egg cell functions.

### 4.4 The maintenance, not initiation, of synergid specific genes were defective in *myb98*

Previously, it has been reported that *MYB98pro∷GFP* is expressed in all cells of the female gametophyte, except for the antipodal cells at FG5 (Ingouff et al., 2006). However, the *MYB98pro∷GFP* signals were detected before the third mitosis, i.e., before cellularization (Figure 3B). Therefore, it is considered that the expression was observed in all cells of the female gametophyte, not only in the synergid cells at FG5. In the *MYB98pro∷NLS–mRuby2*, the fluorescent signals of the NLS–mRuby2 were also detected in all of the female gametophyte cells at FG5 (Figure 4A). Except for the synergid cells, the fluorescent signals of the GFP and NLS–mRuby2 were decreased as the cells matured (Figure 3B, 4A). These results suggested that the synergid cell fate stabilized the gene expression of the *MYB98*. This stabilization was independent of the MYB98. The ectopic expressions of the *MYB98pro∷GFP* and *MYB98pro∷NLS–mRuby2* were not detected after the restrictions of the expression to the synergid cells in the *myb98* mutant (Figure 4B). This suggested that the egg and central cells regularly maintain their cell fates, and the initiation of the synergid cell fate was normal in the *myb98*. Considering these results, the positional information of nuclei is essential for the initiation of the synergid cell fate. Recently, Zhang et al. (2020) reported that AGL80 directly represses the MYB98 expression in the central cell. The specific genes for the accessory cells are expressed in the central cell of *agl61* mutant and *agl80* mutant (Steffen et al., 2008; Zhang et al., 2020). In the egg cell and antipodal cells, the MYB98 expression may also be suppressed by unknown factors. The signal of *SBT4.13pro∷SBT4.13–mClover* was also detected in the synergid cell and the antipodal cells of the *myb98* mutant (Figure 8B). This ectopic expression coincided with the *SBT4.13pro∷SBT4.13–mClover* expression in the egg cell. These findings suggested that the MYB98 may also play a role in preventing the acquisition of the egg cell fate in the accessory cells.

### 4.5 Cell-cell communication between the two synergid cells

An interesting phenotype of the *myb98* mutant was that one of the two synergid cells tends to be converted to an egg cell fate (Table 2; 89% for *CDR1L2–mClover*, 74% for *SBT4.13pro∷SBT4.13–mClover*). Some mutants show similar phenotypes with additional egg cells (Groß-Hardt et al., 2007; Moll et al., 2008; Pagnussat et al., 2007; Kirioukhova et al., 2011; Kong et al., 2015). In the *amp1* mutant, 19% of the ovules showed *EC1.1pro∷HTA6–3GFP* expression in both synergid cells, whereas 26% of the ovules showed their expression in only one of the synergid cells (notably, 45% of the ovules have no detectable fluorescent signal) (Kong et al., 2015). The synergid cells play an important role in the pollen tube attraction, by secreting peptides (Mizuta and Higashiyama, 2018). Previously, we found that the laser disruption of the immature egg cells affects the cell differentiation for one of the synergid cells in the *Torenia fournieri* (Susaki et al., 2015). The results presented here suggest that not only is there cell–cell communication between the egg and synergid cells, but also that there is cell–cell communication between the two synergid cells. Based on these findings, we speculate that the synergid cells detect the abnormal conditions of the egg cell, inducing the decrease of *MYB98* expression. The combination of this monitoring system and the flexible fate maintenance might allow for only one of the two synergid cells to become an egg cell. In the case of the ovule, which has been converted from both the synergid cells to the egg cell fate, it cannot attract pollen tubes. Therefore, it is expected that plants may have a mechanism, which is independent of the MYB98, to retain not only the egg cell but also the synergid cell for pollen tube attraction and fertilization.

Our results suggested that the cell fate specification are immediately initiated around the time of cellularization, depending on the positional information of the nucleus. Moreover, the failure of the cell fate maintenance, like that of the *myb98* mutant, induced cell fate conversions from the adjacent accessory cells to the gametes for compensation of the fertilization. Previously, the existence of the cell–cell communication between the gametic cells and accessory cells, such as lateral inhibition from the egg cell to the synergid cells, was proposed (Tekleyohans et al., 2017). We proposed that the synergid cells communicated with each other to determine their fate and behavior, and such flexibility compensates for the robustness of plant fertilization. Further studies, such as single-cell transcriptome profiling of the mutant synergids, will provide novel insights into the molecular mechanisms of the cell–cell communications in the cell fate specification of plants.

## Author Contributions

D.S. and D.K. conceived the study; D.S. and D.K. designed the experiments; D.S. and T.H. carried out transcriptome analysis; T.S. performed RNA sequencing; D.S. and T.S. analyzed the sequencing data; D.S. and D.M. carried out SBT4.13 expression analysis; M.U. carried out observation of *myb98* embryos; D.K. carried out live-imaging analysis; D.S. and D.K. analyzed the data; D.S., T.H. and D.K. supervised the project; D.S. and D.K. drafted the manuscript; and T.S., D.M., M.U. and T.H. edited the manuscript.

## Acknowledgments

We thank R. Groß-Hardt, T. Kinoshita, and M. Fujimoto for the plant materials; S. Nasu, T. Nishii, T. Shinagawa, and Y. Taniuchi for assistance with cloning and the generation of transgenic plants; T. Nagata, N. Kurata, J. Kawarama, H. Ohyanagi, A. Toyoda, A. Fujiyama for support to examine the method of mRNA amplicfication; H. Nagata and K. Tonosaki for discussions regarding the data analysis; Editage (www.editage.com) for English language editing. Computations were partially performed on the NIG supercomputer at ROIS National Institute of Genetics. This pre-print manuscript was written in LATEXusing a custom style provided by Alex S. Baldwin (github.com/alexsbaldwin/biorxiv-inspired-latex-style) with some modifications.

## Supplemental Data

The supplemental materials are available in the online version of this article.

## Data Availability Statement

RNA-seq data associated with this study have been deposited in DDBJ Sequence Read Archive (DRA) under the accession number, DRR220104–DRR220111. Public data of egg cell, ovule and seedling were DRR174980, DRR174981, DRR174982, DRR044370, DRR066525, SRR346552, SRR346553.

