## Supplemental file for "Dynamics of the cell fate specifications during female gametophyte development in *Arabidopsis*"

### Supplemental Data

#### Supplemental Movies

**Movie S1. Nuclei dynamics in the female gametophyte development in *Arabidopsis thaliana*.**
Time-lapse movies of ovules of *GPR1pro::H2B–mNeonGreen*. Images were taken at 5-min intervals and the movie is displayed at 30 frames per second. Scale bar, 20 $\mu$m (see also Figure 1B).

**Movie S2. Morphological changes and plasma membrane formation in the female gametophyte development**
Time-lapse movies of ovules of *RPS5Apro::tdTomato–LTI6b*. Images were taken at 5-min intervals and the movie is displayed at 30 frames per second. Scale bar, 20 $\mu$m (see also Figure 2A).

**Movie S3. Nuclei dynamics during plasma membrane formation**
Time-lapse movies of ovules of *RPS5Apro::H2B-sGFP* (green) and *RPS5Apro::tdTomato–LTI6b* (magenta). Images were taken at 10-min intervals and the movie is displayed at 15 frames per second. Scale bar, 20 $\mu$m (see also Figure 2C).

**Movie S4. The maturation of female gametophyte cells after cellularization**
Time-lapse movies of ovules of *RPS5Apro::tdTomato–LTI6b*. Images were taken at 5-min intervals and the movie is displayed at 30 frames per second. Scale bar, 20 $\mu$m (see also Figure 2D).

**Movie S5. Expression of egg cell–specific markers at FG5**
Time-lapse movies of ovules of *EC1.2pro::mtKaede* (green) and *ABI4pro::H2B–tdTomato* (magenta). Images were taken at 10-min intervals and the movie is displayed at 15 frames per second. Scale bar, 20 $\mu$m (see also Figure 3A).

**Movie S6. Expression of synergid cell–specific marker at FG4 and FG5**
Time-lapse movies of ovules of *MYB98pro::GFP* (green) and *RPS5Apro::H2B–tdTomato* (magenta). Images were taken at 10-min intervals and the movie is displayed at 15 frames per second. Scale bar, 20 $\mu$m (see also Figure 3B).

**Movie S7. Expression of egg cell and synergid cell–specific markers after cellularization**
Time-lapse movies of ovules of *RPS5Apro::tdTomato–LTI6b* (magenta), *EC1.1pro::NLS–3xDsRed* (magenta), and *LURE1.2pro::NLS–3xGFP* (green). Images were taken at 5-min intervals and the movie is displayed at 30 frames per second. Scale bar, 20 $\mu$m (see also Figure 3C).

**Movie S8. Expression of egg cell, synergid cell, and antipodal cell–specific markers after cellularization**
Time-lapse movies of ovules of *EC1.1pro::SP-mTurquoise2–CTPP* (cyan), *MYB98pro::mRuby3–LTI6b* (magenta), *DD1pro::ermTFP1* (green), and *AKVpro::H2B–mScarlet-I* (magenta). Images were taken at 10-min intervals and the movie is displayed at 15 frames per second. Scale bar, 20 $\mu$m (see also Figure 3E).

**Movie S9. Expression of *MYB98pro::NLS-mRuby2* in wild type**

Time-lapse movies of ovules of *MYB98pro::NLS–mRuby2* in wild type. Images were taken at 10-min intervals and the movie is displayed at 15 frames per second. Scale bar, 20 $\mu$m (see also Figure 4A).

**Movie S10. Expression of *MYB98pro::NLS-mRuby2* in *myb98***

Time-lapse movies of ovules of *MYB98pro::NLS–mRuby2* (magenta) and *MYB98pro::GFP* (green) in *myb98*. Images were taken at 10-min intervals and the movie is displayed at 15 frames per second. Scale bar, 20 $\mu$m (see also Figure 4B).

**Movie S11. Expressions of CDR1s–mClover after cellularization**

Time-lapse movies of ovules of *CDR1–LIKE2pro::CDR1–LIKE2–mClover* (green) in the first movie, *CDR1–LIKE1pro::CDR1–LIKE1–mClover* (green) in the second movie, and *CDR1pro::CDR1–mClover* (green) in the third movie. Images were taken at 15-min intervals and the movie is displayed at 15 frames per second. Scale bar, 20 $\mu$m (see also Figure 7B).

**Movie S12. Expression of *CDR1–LIKE2pro::CDR1–LIKE2–mClover* in *myb98***

Time-lapse movies of ovules of *CDR1–LIKE2pro::CDR1–LIKE2–mClover* in *myb98*. Images were taken at 10-min intervals and the movie is displayed at 15 frames per second. Scale bar, 20 $\mu$m (see also Figure 7C).

**Movie S13. Expression of *SBT4.13pro::SBT4.13–Clover* in wild type**

Time-lapse movies of ovules of *SBT4.13pro::SBT4.13–mClover* in wild type. Images were taken at 10-min intervals and the movie is displayed at 15 frames per second. Scale bar, 20 $\mu$m (see also Figure 8A).

**Movie S14. Expression of *SBT4.13pro::SBT4.13–Clover* in *myb98***

Time-lapse movies of ovules of *SBT4.13pro::SBT4.13–Clover* in *myb98*. Images were taken at 10-min intervals and the movie is displayed at 15 frames per second. Scale bar, 20 $\mu$m (see also Figure 8B).

**Movie S15. Expression of egg cell, synergid cell, and antipodal cell–specific markers in *myb98***

Time-lapse movies of ovules of *EC1.1pro::SP-mTurquoise2–CTPP* (cyan), *MYB98pro::mRuby3–LTI6b* (magenta), *DD1pro::ermTFP1* (green) in *myb98*. Images were taken at 10-min intervals and the movie is displayed at 15 frames per second. Scale bar, 20 $\mu$m (see also Figure 8C).

#### Supplemental Tables

**Table S1. Frequency of available synergid cells using enzyme solution with or without calcium nitrate.**

After an hour enzyme treatment with or without calcium nitrate, we counted the ovules and collectable synergid cell protoplasts which maintained GFP signal and detached from ovule and other cells.

**Table S2. Frequency of collectable synergid cells using enzyme solution (pH 5–9).**

We counted the ovules and collectable synergid cell protoplasts which maintained GFP signal for an hour enzyme treatment (pH 5–9).

**Table S3. TPM of all expression genes and stastistics for DEGs identificatiion.**

Transcripts per million (TPM) value of the genes (raw read > 10) and statistical processing for DEGs identification among egg, cenrtral and synergid cells or between synergid cells in wild type and *myb98*.

**Table S4. Egg specific genes and DEGs between wild type and *myb98* synergid cells.**

The genes specifically expressed in the egg cells that were DEGs between wild type and *myb98* synergid cells.

#### Supplemental Figure

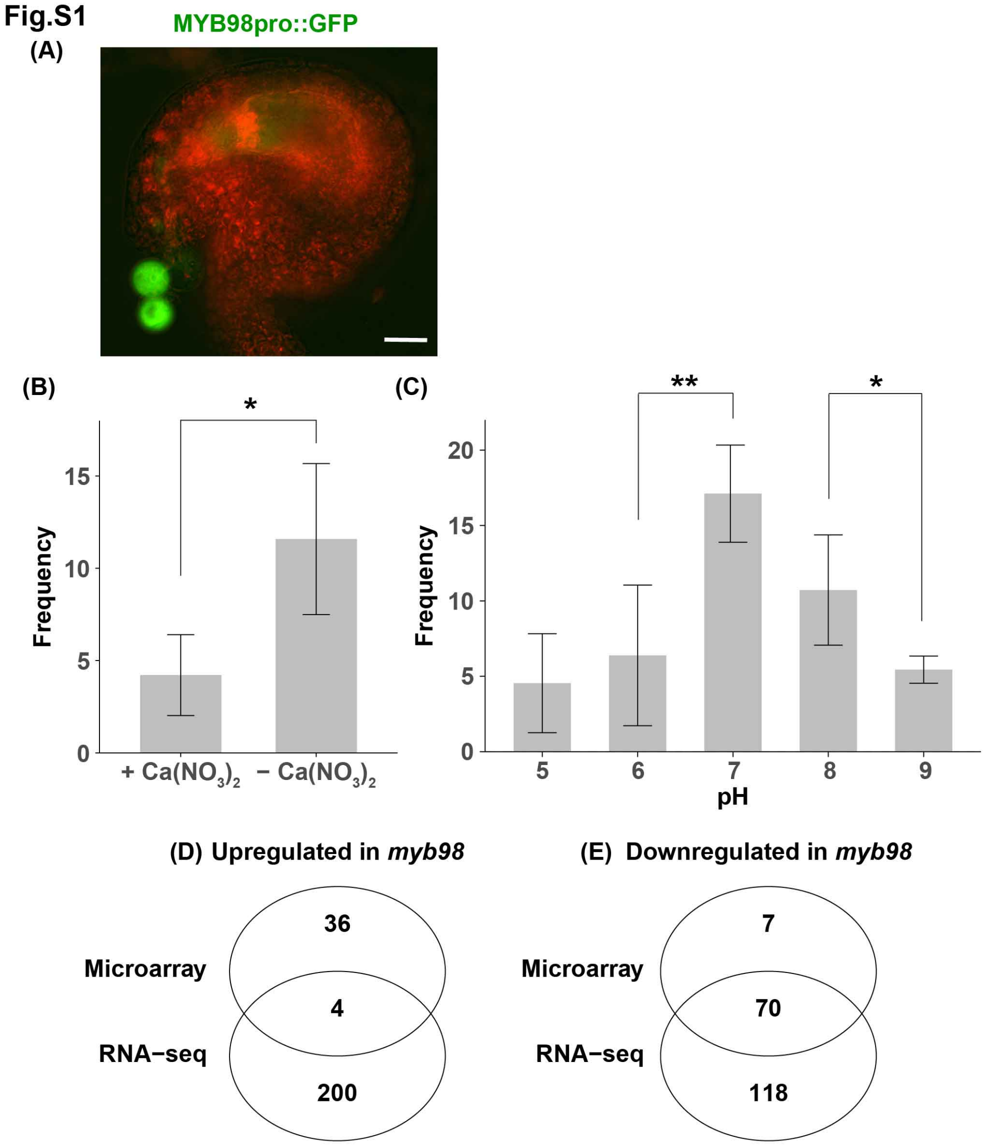


Figure S1: RNA-seq of the female gametophyte cells. **(A)** Two synergid cells were released from the ovules of *MYB98pro::GFP*. **(B)** Frequency of the collectable synergid cells with or without calcium nitrate in the enzyme solution. F test of frequency showed that there is no significant difference between the two variances ($p > 0.05$). Results of the t-test showed that the absense of calcium nitrate was more effective for synergid cell isolation (*$p < 0.05$). **(C)** Frequency of the collectable synergid cells depending on the pH of the enzyme solution. F test of frequency showed that there is no significant difference in variances between pH 6 and pH 7 or pH 7 and pH 8 ($p > 0.05$). Results of the t-test showed that the frequency of collectable synergid cells was different significantly between pH 6 and pH 7 or pH 7 and pH 8 (*$p < 0.05$; **$p < 0.01$). **(C, D)** Venn diagram of differentially expressed genes upregulated **(D)** or downregulated **(E)** in the *myb98* mutant synergids between RNA-seq and microarray.
